# McClintock: An integrated pipeline for detecting transposable element insertions in whole genome shotgun sequencing data

**DOI:** 10.1101/095372

**Authors:** Michael G. Nelson, Raquel S. Linheiro, Casey M. Bergman

## Abstract

**Background:** Transposable element (TE) insertions are among the most challenging type of variants to detect in genomic data because of their repetitive nature and complex mechanisms of replication. Nevertheless, the recent availability of large resequencing datasets has spurred the development of many new methods to detect TE insertions in whole genome shotgun sequences. These methods generate output in diverse formats and have a large number of software and data dependencies, making their comparative evaluation challenging for potential users.

**Results:** Here we develop an integrated bioinformatics pipeline for the detection of TE insertions in whole genome shotgun data, called McClintock (https://github.com/bergmanlab/mcclintock), that automatically runs and generates standardized output for multiple TE detection methods. We demonstrate the utility of the McClintock system by performing comparative evaluation of six TE detection methods using simulated and real genome data from the model microbal eukaryote, *Saccharomyces cerevisiae*. We find substantial variation among McClintock component methods in their ability to detect non-reference insertions in the yeast genome, but show that non-reference TEs at nearly all biologically-realistic locations can be detected in simulated data by combining multiple methods that use split-read and read-pair evidence. In general, our results reveal that split-read methods detect fewer non-reference TE insertions than read-pair methods, but generally have much higher positional accuracy. Analysis of a large sample of real yeast genomes reveals that most, but not all, McClintock component methods can recover known aspects of TE biology in yeast such as the transpositional activity status of families, tRNA gene target preferences, and target site duplication structure, albeit with varying levels of positional accuracy.

**Conclusions:** Our results suggest that no single TE detection method currently provides comprehensive detection of non-reference TEs, even in the context of a simplified model eukaryotic genome like *S. cerevisiae*. In spite of these limitations, the McClintock system provides a framework for testing, developing and integrating results from multiple TE detection methods to achieve this ultimate aim, as well as useful guidance for yeast researchers to select appropriate TE detection tools.

## Background

The widespread availability of genomic data over the last two decades has provided unparalleled opportunities to learn about the abundance, diversity, and functional consequences of transposable elements (TEs) in modern genomes. However, the computation analysis of TE sequences in both reference and resequenced genomes remains a challenging area of bioinformatics research because of the repetitive nature of these sequences. Development of bioinformatics tools for the detection and annotation of TEs in reference genomes is now a relatively mature field [1–3], although many open questions remain about choosing the best tools for specific biological applications [4]. In contrast, detection of reference and non-reference TE insertions in whole-genome shotgun (WGS) resequencing data is an active research area (reviewed in [5]), with a wide array of methods published in recent years [6–32].

Because of the wide array of available methods, it remains unclear which method for detecting TEs in resequenced genomes is best suited for particular genomic problems, leading to substantial investigator effort in terms of installation and testing, or the application of sub-optimal bioinformatic approaches. Most papers reporting new methods to detect reference or non-reference TEs in WGS data provide some measure of their own performance to relative using simulations, benchmark genomic data, or PCR-based validation. However, only a handful of papers have reported new methods that include performance evaluation relative to other methods [5, 23, 25, 27–29, 32], and these are often limited in scope to only a single organism or TE family. In addition to being incomplete, comparative analysis of bioinformatic systems in papers that report new methods can fall victim to the “self-assessment trap” [33]. Moreover, there is no common format for the annotation of non-reference TE insertions [34, 35], making direct comparison of predictions from different methods more challenging. Recently, Rishishwar *et al.* [35] performed an independent comparative evaluation of seven WGS-based TE detection methods using human genomic data, which revealed many method-specific predictions and recommended combining the results of multiple systems followed by manual curation (see also [5]). Rishishwar *et al.* [35] also highlighted the challenges users face when installing and running multiple TE detection methods, and provide helpful advice for users and developers.

As a step towards a fully automated framework for running and evaluating multiple methods to detect TEs in WGS resequencing data, we have developed an integrated pipeline called McClintock (https://github.com/bergmanlab/mcclintock) that generates standardized output for multiple WGS-based TE detection methods. The primary goal of the McClintock pipeline is to lower the barrier to installation, use, and evaluation of multiple WGS-based TE detection methods. Several key features of the McClintock pipeline are that it automates formatting of key input files and standardizes output of multiple TE detection methods to allow easy comparisons of results from different systems, as recommended by [35]. In the initial version of McClintock, we incorporate six complementary TE detection methods that make predictions based on split-read or read-pair based evidence in Illumina WGS data. Here we describe the McClintock system and its component methods, and perform comparative evaluation using simulated and real yeast genome data. Our analysis supports previous conclusions that no single TE detection method provides comprehensive detection of non-reference TEs [5, 35], but provides a framework for further testing, development and integration to achieve this ultimate aim, as well as useful guidance for yeast researchers to select appropriate TE detection tools.

## Implementation

### McClintock component methods and their dependencies

We initiated our design of McClintock with a literature search for candidate bioinformatic systems that can detect TE insertions from NGS data in 2014, which yielded 33 potential systems. Our main project objective was to develop a system that automatically detects non-reference TE insertions in raw WGS data for any species. Thus, we excluded systems that required any wet-lab enrichment further consideration. Systems that did not make their code available were also rejected. This left a list of 12 candidate software systems. After preliminary testing of these 12 methods, six were rejected from further testing because of difficulties during installation (Tangram [21], VariationHunter [9]), reliance on data for a specific organism (TEA [13], VirusSeq [18]), inability to detect non-reference insertions (T-lex [11]), or the inability to distinguish general structural variations from TE insertions (HYDRA [10]). Six remaining methods (ngs_te_mapper [12], TE-locate [14], PoPoolationTE [15], RetroSeq [36], RelocaTE [17] and TEMP [20]) had publicly-available code that could be installed reproducibly and met project objectives were selected for incorporation into the initial McClintock pipeline. Since the original selection of methods for inclusion in McClintock, a number of additional methods that meet the initial project requirements (“pecnv teclust” [19], TIF [22], TE-Tracker [23], Mobster [24], ITIS [25], Jitterbug [27], TIDAL [28], ISmapper [29], MELT [37] SPLITREADER [30], and TEPID [38]) and new versions of some methods (PoPoolationTE2 [31] and RelocaTE2 [32]) have been released. These methods have not yet been incorporated into McClintock, but the flexible architecture of the system permits their inclusion in the future.

A summary of the main features of the six component methods included in McClintock is shown in Table 1. These six component systems have many dependencies on other pieces of software, which must all be correctly installed before the component system will function correctly. These software dependencies are listed in Table 2. Several of these component dependencies require end user licenses, and thus it was not possible to fully automate installation of all component methods. McClintock therefore assumes component dependencies are installed system-wide, but automates installation of the component methods themselves. A passive check is performed during installation of McClintock that reports whether component dependencies are available, though installation is not halted if they are missing. Because of the large number of component dependencies and subsequent development of components themselves, we developed McClintock to use specific versions of components and their dependencies. Table 2 also lists the version of each dependency that was used with McClintock to obtain the results presented here.

**Table 1:** An overview of the features of the component TE detection methods in the McClintock pipeline. Split-read and read-pair refer to what type of evidence is used to make TE insertion predictions (see Implementation in Main Text for details).

| Method | ngs_te_mapper | RelocaTE | TEMP | RetroSeq | PoPoolationTE | TE-locate |
| --- | --- | --- | --- | --- | --- | --- |
| Split-read | ✓ | ✓ | ✓ |  |  |  |
| Read-pair |  |  | ✓ | ✓ | ✓ | ✓ |
| Non-reference TEs | ✓ | ✓ | ✓ | ✓ | ✓ | ✓ |
| Reference TEs | ✓ | ✓ | ✓<sup>i</sup> |  | ✓ | ✓ |
| Orientation | ✓ | ✓<sup>ii</sup> | ✓ |  |  | ✓<sup>iii</sup> |
| TSD | ✓ | ✓ | ✓<sup>iv</sup> |  |  |  |
| Detects TE families not in reference genome | ✓ | ✓ | ✓ | ✓<sup>v</sup> | ✓ |  |

<sup>i</sup> TEMP reports whether a reference TE is absent from the resequenced sample rather than providing direct evidence for the presence of a reference TE. <sup>ii</sup> RelocaTE output provides information about the orientation of non-reference TEs, but not for reference TEs. McClintock annotates the orientation of reference TEs in RelocaTE output using the original reference TE annotation. <sup>iii</sup> TE-locate provides information about the orientation of non-reference TEs where possible, but not for reference TEs. McClintock annotates the orientation of reference TEs in TE-locate output using the original reference TE annotation. <sup>iv</sup> TEMP only makes TSD predictions for insertions with split-read support. <sup>v</sup> RetroSeq can detect TE families not present in the reference genome when using Exonerate to generate a reference TE annotation, but not when using a user-supplied reference TE annotation that is the default option in McClintock.

**Table 2:** Software dependencies required to install and run each component TE detection method in the McClintock pipeline.

| Software | ngs_te_mapper | RelocaTE | TEMP | RetroSeq | PoPoolationTE | TE-locate | Version used in this study |
| --- | --- | --- | --- | --- | --- | --- | --- |
| Linux | ✓ | ✓ | ✓ | ✓ | ✓ | ✓ | CentOS 6 |
| Perl |  | ✓ | ✓ | ✓ | ✓ | ✓ | 5.18.1 |
| R [64] | ✓ |  |  |  |  |  | 3.0.2 |
| BioPerl [65] |  | ✓ | ✓ |  |  |  | 1.006001 |
| RepeatMasker [66] |  |  |  |  | ✓ |  | 4.0.2 |
| BEDTools [60] |  |  | ✓ | ✓ |  |  | 2.17.0 |
| SAMTools [58] |  | ✓ | ✓<sup>i</sup> | ✓ | ✓ |  | 0.1.19-44428cd |
| BCFTools [58] |  |  |  | ✓ |  |  | 0.1.19-44428cd |
| twoBitToFa [67] |  |  | ✓ |  |  |  | 294 |
| BLAT [68] |  | ✓ |  |  |  |  | 35x1 |
| Exonerate [69] |  |  |  | ✓ |  |  | 2.2.0 |
| Bowtie [70] |  | ✓ |  |  |  |  | 1.0.0 |
| BWA [71] | ✓ |  | ✓ | ✓ | ✓ | ✓ | 0.7.4-r385<sup>ii</sup> |

<sup>i</sup> Only compatible with SAMTools 0.1.19 or earlier [35].<sup>ii</sup> This specific version of BWA is needed to ensure compatibility between PoPoolationTE, which uses BWA-ALN, and other component methods that use BWA-MEM.

McClintock component methods also have a variety of data dependencies that are required as inputs, which are listed in Table 3. The component methods incorporated into McClintock together require a total of 13 different data dependencies to run. However, since many of these data dependencies can be automatically generated or are format alterations that can be automatically achieved with simple pre-processing steps, the number of data dependencies can be reduced to three required inputs for McClintock: a fasta file of the reference genome, a fasta file of the canonical TE sequences, and fastq files of NGS reads (paired or single ended).

**Table 3:** Data dependencies required to successfully run each component of the McClintock pipeline

|  | ngs_te_mapper | RelocaTE | TEMP | RetroSeq | PoPoolationTE | TE-locate |
| --- | --- | --- | --- | --- | --- | --- |
| Reference genome (fasta) | ✓ | ✓ | ✓ | ✓ | ✓ | ✓ |
| Canonical TE sequences (fasta) | ✓ | ✓<sup>i</sup> | ✓ | ✓<sup>ii</sup> | ✓ |  |
| Annotation of reference TEs (GFF) |  |  |  |  | ✓ | ✓ |
| Annotation of reference TEs (BED) |  |  | ✓ | ✓<sup>iii</sup> |  |  |
| Annotation of reference TEs (custom format) |  | ✓ |  |  |  |  |
| Unaligned reads (single-end fastq) | ✓ | ✓ |  |  |  |  |
| Unaligned reads (paired-end fastq) |  |  |  |  | ✓ |  |
| Aligned reads (BAM) |  |  | ✓ | ✓ |  |  |
| Aligned reads (lexically sorted SAM) |  |  |  |  |  | ✓ |
| TE hierarchy (custom format) |  |  | ✓ |  | ✓ |  |

<sup>i</sup> Must include an entry in the format “TSD=...” for each TE in the file on the same line as the header, where “...” is the TSD sequence if known, or a string of periods with equal to the TSD length if the TSD sequence is unknown. If neither length nor the sequence of the TSD is known, “TSD=UNK” can be supplied. <sup>ii</sup> Must be formatted as one fasta file per TE family and a file-of-files listing their locations. <sup>iii</sup> Must be one BED file for each entry in the reference TE annotation and a file-of-files listing their locations.

A more detailed overview of the component methods, their software/data dependencies, and limitations is provided in the Description of McClintock Component Methods section of Additional File 1.

### The McClintock Pipeline

An overview of the data flow and processing steps performed by the McClintock pipeline is shown in Figure 1. In the following sections, we describe the options for running the McClintock pipeline, then describe how component methods are executed and parsed in the context of the McClintock pipeline.

**Figure 1.**
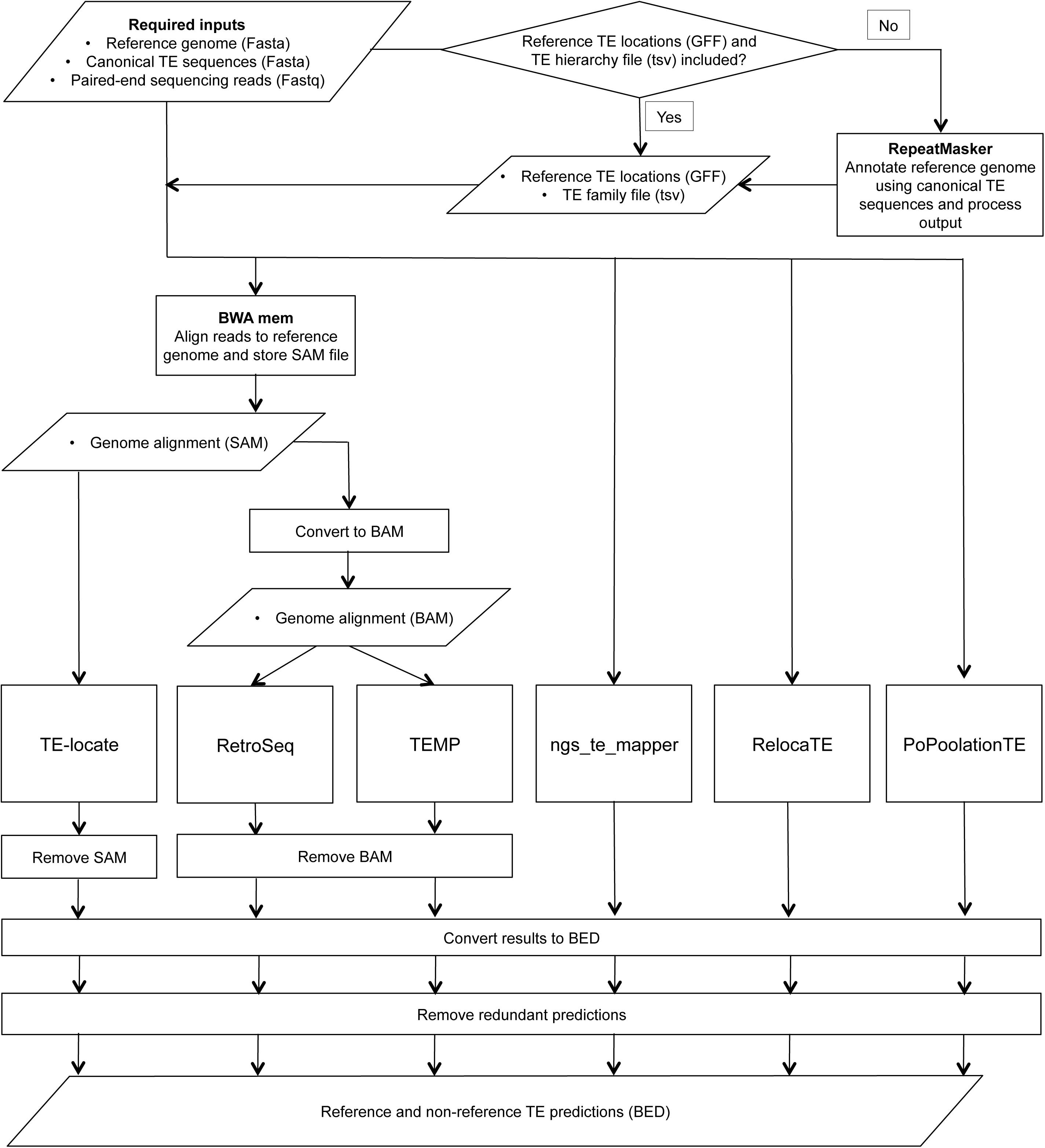
Overview of the McClintock pipeline. The flowchart shows important processes are shown as boxes, decision points as diamonds, and data at important steps as parallelograms. Note that the last three steps of the pipeline are applied independently to each method. Final results from each component method are output independently by McClintock, allowing the user to easily merge output or assess for overlap among methods.

#### Options

##### Reference TE annotation options

If a pre-existing annotation of the TE sequences in the reference genome is available, a one-based GFF file of this data can be used as input for the McClintock pipeline. If a reference TE annotation is provided, then the user must also create and supply a TE “hierarchy” file as another input. The hierarchy file contains two tab-delimited columns, the first listing the name of each instance in the reference TE annotation and the second listing the canonical TE family that instance belongs to. If no reference TE annotation is provided, then a reference TE annotation and hierarchy file is created automatically by running RepeatMasker and post-processing RepeatMasker output files.

##### Reference genome sequence options

McClintock provides options to automatically create various different modified reference genomes. These options were implemented because some component methods (RetroSeq and TE-locate) require an instance of a TE to exist in the reference genome for non-reference instances of that family to be detected in a resequenced sample. This is important because, in some cases, like the *D. melanogaster P*-element [39], the reference genome does not include any copies of a TE family that occurs in natural populations. This situation may also occur when a TE family has been introduced experimentally into a strain lacking that TE to study its transposition. To allow for these cases, McClintock has an option to generate modified reference genomes that include additional “chromosomes” comprised of canonical TE sequences or TE sequences extracted from the reference genome. An annotation of TEs in the additional “chromosomes” is then appended to the reference TE annotation file. PoPoolationTE requires a modified reference genome with canonical TE sequences and reference TE sequences added as additional “chromosomes.” Thus these reference genome modifications are always made specifically for PoPoolationTE, regardless of whether user-supplied options to modify the reference genome are provided globally for other component methods.

##### Run options

McClintock offers additional options to customize the way the pipeline is run. It is possible to specify which component methods are executed, allowing tailored output and shorter run times. McClintock and its component methods produce short-read alignment files and other intermediate files that can be very large, and thus an option is provided to remove unwanted intermediate files. BAM files output by McClintock may be useful for other purposes, so an option is provided to eliminate all intermediate files other than BAM files. The location of all output files can be changed to any absolute path that the user requests. Within the specified location, all output files will be produced in a directory named after the reference genome sequence with results for each sample stored in subdirectories named after the fastq files for that sample, allowing multiple samples for the same reference genome to re-use common index files.

#### Overview of the McClintock process

From the limited set of inputs and options the user provides, McClintock then automatically generates all other input files required to run all six component methods. If the reference TE annotation and hierarchy file are provided by the user, the RepeatMasker step is skipped and the user-supplied reference TE annotation is used to make a hard-masked version of the reference genome using BEDTools (a step that is required only for PoPoolationTE). If the reference TE annotation and TE hierarchy file are not supplied by the user, McClintock launches RepeatMasker, which creates a reference TE annotation in GFF format that is in turn used by McClintock to create the TE hierarchy file. If specified, modifications can be made to the reference genome prior to automatic generation of the reference TE annotation and hierarchy file (see Options section above). McClintock then converts the reference TE annotation to BED format, as required by TEMP and RetroSeq.

Prior to running any of the component methods, McClintock runs FastQC on the input fastq files to provide the user information to help interpret McClintock output. FastQC results are stored in a quality control subdirectory for each sample. Next, all indexing steps for the reference genome are performed. If only single-ended NGS data is provided, this is automatically detected by McClintock, and only the component methods that can analyse single-ended NGS data (ngs_te_mapper and RelocaTE) are launched. In this case, the main BWA-MEM alignment step is not performed because ngs_te_mapper and RelocaTE execute their own internal alignments. If paired-end NGS data is provided, then the main BWA-MEM alignment of the NGS data to the reference genome is launched and stored in SAM format. If TE-locate or TEMP are to be run, then the median insert size is calculated based on the distance between aligned pairs of reads in this SAM file. If TE-locate is to be run, then the SAM output of BWA-MEM is lexically sorted and a new SAM file is retained. If TEMP or RetroSeq are to be launched, then the SAM alignment file is sorted, converted into BAM format and indexed. In addition, if a BAM file is created, then McClintock will launch SAMtools flagstat to produce mapping summary statistics that are stored in the quality control subdirectory for each sample.

To launch ngs_te_mapper, the basic inputs to McClintock are sufficient and no additional pre-processing is required. To launch RelocaTE, “TSD=UNK” is automatically added to each identifier line in the canonical TE fasta file, providing maximum flexibility for this method. The custom reference TE annotation required by RelocaTE is produced from the user-supplied GFF or created from RepeatMasker output. To run TEMP, soft links are created to the BAM and BAM index files to ensure they have the required suffixes (“sorted.bam” and “sorted.bam.bai,” respectively). To run RetroSeq, the canonical TE file or reference TE annotation file is split into one file per TE family, and a file-of-files is produced with these file locations. For RetroSeq, McClintock uses the less computationally-intensive approach of assigning discordant reads to a TE family based on reference TE locations, rather than alignment of the reads to canonical TE sequences using Exonerate (we found the latter approach caused frequent failures during testing). The code to run the Exonerate step is included in McClintock if a user has data and a compatible computing environment. To launch PoPoolationTE, the basic TE hierarchy file is reformatted to add additional columns required by this method. Also, the identifiers of reads in the fastq input files are also changed so that they end with “\1” or “\2” for each member of a pair of reads. Finally, the median insert size of fragments is calculated based on the distance between aligned pairs of reads in a PoPoolationTE-specific SAM file (created using the BWA-ALN algorithm), and the read length is obtained from the fastq files. These values are passed to a patched version of PoPoolationTE that allows sample-specific parameters to be set for clustering TE-supporting reads. To run TE-locate, the reference TE annotation file is modified using the TE hierarchy file to ensure that the correct family level of annotation is provided in the column required by TE-locate. TE-locate also requires that the reference genome has more than five chromosomes. Should this not be the case, McClintock will add as many false chromosomes as required to produce five in total. Once these pre-processing steps are performed, each of the component methods are run following the guidelines described in their publications and manuals. (See Description of McClintock Component Methods in Additional File 1 for further details).

To make McClintock runs more efficient for large resequencing datasets from the same species, input files that are reference genome specific but not sample specific (for example, genome indexes and reference TE annotations) are saved separately in the highest level of the output directory. If another sample is run for the same reference genome in the same output location, then these files can be reused, saving both space and time. As noted above, files that are not required, such as intermediate output and large genome alignments, can be deleted once used to minimize disk space held throughout the run. Also, if a subcomponent of McClintock is not run then, where possible, McClintock will not create any input files that are solely required for that method.

#### Post-processing and standardization of output format

The component methods within McClintock produce their output in different file formats and annotation frameworks (see [34] for discussion). Therefore, McClintock performs a number of post-processing steps to standardize outputs from different methods into a common annotation framework. Details of the native annotation framework for component methods and the post-processing steps made by McClintock can be found in the Post-processing and Standardisation of Component Method Output section of Additional File 1). Before performing these steps, the original (unedited) results for each method are saved in the output directory for that sample. If TE predictions are made by any component method in the additional “chromosomes” added in modified reference genomes (see Options section above), these results are removed from the standard results files and retained in a subdirectory within the results directory called “non-ref_chromosome_results”.

The output file format chosen to standardize results for all component methods is zero-based BED6 format, because it allows easy integration with the BEDTools and UCSC genome browser. BED format provides a fourth column to contain a name for the annotated feature. All records in these BED files contain the name of the TE family predicted at that location and whether the prediction is of a non-reference or reference TE. The name column also reports the sample ID from the fastq input file and the name of the component method that made the prediction. The type of evidence used for the prediction is also listed, either “sr” representing a prediction made from split-read evidence, “rp” representing a prediction made from read-pair evidence, or “nonab” for TEMP reference TE predictions that rely on no evidence for the absence of the TE in the sample. In addition, filtering and redundancy removal was performed within the result file for each component method. No redundancy filtering is performed by McClintock across component methods, allowing users to more directly compare output from different methods. To facilitate viewing of results on the UCSC genome browser, a header is included in each BED file. This header is read by the UCSC browser and lists the sample name and McClintock component system that produced the results as the track name and description, allowing multiple result files for the same sample to be merged and visualized simultaneously.

## Results and Discussion

### Application of McClintock to simulated *S. cerevisiae* genomes with single synthetic TE insertions

To test McClintock and its component methods, we used simulated WGS datasets based on the genome of the model eukaryote, *S. cerevisiae*. We chose *S. cerevisiae* for testing McClintock because its reference genome is relatively small and has been completely determined [40], it has large samples of publicly-available resequenced genomes [41–43], and the genome biology of its TEs is well-characterized [44, 45]. In addition, the six TE families in *S. cerevisiae* (*Ty1*, *Ty2*, *Ty3*, *Ty3*_*1p*, *Ty4*, and *Ty5*) are all long terminal repeat (LTR) retrotransposons, a type of TE that can be processed effectively by all six McClintock component methods. We first performed control analyses by simulating WGS resequencing of unmodified *S. cerevisiae* reference genome samples and applying McClintock to these datasets (see Simulating Resequencing of the *S. cerevisiae* Reference Genome in Additional File 1). While not the major focus of this study, these reference genome simulations allowed us to evaluate how often McClintock component methods detected reference TEs and, more importantly, how often component methods detected false positive non-reference TEs (in the absence of any true, non-reference TE insertions). An example of reference TE predictions for all six component methods is shown in Additional Figure 1A. In general, analysis of unmodified simulated reference genomes showed that McClintock component methods cannot detect all reference TEs, but also typically have low false positive rates for predicting non-reference TE insertions when they are truly absent (see Additional Table 1 and Additional Table 2). Additionally, these simulations showed that McClintock had better performance at 100X versus 10X coverage, and that neither the choice of reference TE annotation nor reference genome options substantially affected the detection of reference or non-reference TEs for most McClintock component methods.

Next, we simulated WGS samples for reference genomes that include a single synthetic TE insertion (placed at biologically-realistic locations upstream of tRNA genes) to evaluate the ability of McClintock component methods to detect true positive non-reference TE insertions. To do this, WGS reads were simulated for 598 samples, each with a different synthetic TE insertion placed upstream of one of the 299 tRNA genes in the yeast genome. 299 samples were created for synthetic insertions the positive orientation upstream of tRNA genes and 299 samples for synthetic insertions in the negative orientation. Genomes with synthetic insertions were created by selecting a five bp sequence 12-17bp upstream of a tRNA start site for *Ty3* or 195-200bp upstream of a tRNA start site for *Ty1*, *Ty2*, and *Ty4*. This five bp sequence formed the basis of a synthetic target site duplication (TSD) and became the location into which a full-length *Ty* canonical sequence was inserted in the sacCer2 reference genome. All single insertion samples were simulated at 100X coverage since the ability of component methods to detect reference TEs improved with increasing coverage and to match analysis of real yeast genomes (see Application of McClintock to 93 yeast genomes). An illustration of non-reference TE predictions for all six component methods in a genomic segment containing a synthetic TE insertion is shown in Additional Figure 1B. In the following sections, we detail the analysis of these single synthetic insertion simulated samples in terms of overall numbers of reference and non-reference TE predictions and positional accuracy of non-reference TE predictions.

#### Numbers of reference and non-reference TE predictions

Table 4 shows the mean number of reference and non-reference TE insertions predicted across all 299 simulated single-insertion samples on the positive and negative strands, respectively. Table 4 also shows the mean number of correct non-reference insertions predicted per sample at a given distance threshold. If all single TE insertion samples were predicted correctly for a method, it would lead to an average value of exactly one non-reference TE predicted per sample. Comparing row one of Table 4 (single insertion simulation) with row nine of Additional Table 1 (unmodified reference simulation), we can infer that the inclusion of single synthetic insertions into the yeast genome does not substantially alter the ability of any McClintock component method to predict reference TEs. As expected, comparing row two of Table 4 (single insertion simulation) with row nine of Additional Table 2 (unmodified reference simulation), we see gains in the numbers of non-reference TE insertions predicted for all methods, demonstrating that McClintock components can detect true positives above false positive baselines in our simulation framework.

**Table 4:** Average numbers of predictions and correct predictions, by method, for simulated yeast WGS samples with a single synthetic TE insertion upstream of tRNA genes. Simulated WGS samples had 100X coverage, and McClintock was run using the reference TE annotation from Carr *et al.* [45] and the unmodified reference genome option. The first two rows show the mean number of reference and non-reference predictions per sample, averaged across all simulated samples for that strand. Rows three to six show the average number of non-reference predictions of the correct TE family across sample that fell within the given distance of the known synthetic TE insertion site. For each method, the first column corresponds to insertions on the positive strand and the second column corresponds to insertions on the negative strand. For a prediction to be considered “exact” the location of the TSD had to be predicted correctly. Numbers for TEMP combine predictions with split-read and read-pair support.

| Insertion strand | ngs.te.mapper |  | RelocaTE |  | TEMP |  | RetroSeq |  | PoPoolationTE |  | TE-locate |  |
| --- | --- | --- | --- | --- | --- | --- | --- | --- | --- | --- | --- | --- |
|  | + | - | + | - | + | - | + | - | + | - | + | - |
| Reference TEs mean | 41.26 | 41.26 | 130.42 | 130.42 | 482.98 | 482.98 | N.A. | N.A. | 163.50 | 163.50 | 271.32 | 271.32 |
| Non-reference TEs mean | 0.42 | 0.32 | 1.12 | 1.11 | 0.90 | 0.90 | 0.87 | 0.86 | 1.18 | 1.14 | 0.98 | 0.92 |
| Exact | 0.40 | 0.29 | 0.30 | 0.24 | 0.36 | 0.36 | 0.00 | 0.00 | 0.00 | 0.00 | 0.00 | 0.00 |
| Within 100bp | 0.40 | 0.29 | 0.63 | 0.61 | 0.90 | 0.90 | 0.68 | 0.66 | 0.16 | 0.16 | 0.07 | 0.06 |
| Within 300bp | 0.40 | 0.29 | 0.63 | 0.61 | 0.90 | 0.90 | 0.69 | 0.67 | 0.16 | 0.16 | 0.70 | 0.54 |
| Within 500bp | 0.40 | 0.29 | 0.63 | 0.61 | 0.90 | 0.90 | 0.69 | 0.67 | 0.16 | 0.16 | 0.82 | 0.78 |

For ngs_te_mapper, the average number of non-reference predictions shows this method systematically under-predicts non-reference TE insertions. However, the average number of predictions made overall per sample is only slightly higher than the average number of exact predictions. Consistent with unmodified reference genome simulations (see row nine of Additional Table 2), this result indicates that only a small number of non-reference predictions made by ngs_te_mapper are false positives. Moreover, whenever ngs_te_mapper makes a prediction of a non-reference TE (that is within 500bp of the true insertion site), the prediction was always at the exact TSD, suggesting high accuracy in terms of position and TSD structure for this method (see below). We also observed that ngs_te_mapper detected fewer insertions when the synthetic insertion is on the negative strand relative to the tRNA gene, suggesting there can be strand bias in the detection of non-reference TEs. This bias could be due to yeast genome organization, our simulation framework, the ngs_te_mapper algorithm or a combination of these factors.

RelocaTE produced, on average, slightly more than one non-reference TE prediction per sample. At face value, this result suggests that RelocaTE may detect essentially every synthetic insertion, but also make occasional false positive predictions. In fact the average excess number of predictions made by RelocaTE in single insertion simulated genomes is very close to the false positive rates observed in simulations of unmodified reference genomes (see row nine of Additional Table 2). However, only about 50% of the total RelocaTE predictions are made within 500bp of the true insertion. Thus, it appears that the inclusion of single synthetic insertions increases the rate of false positive non-reference TE predictions RelocaTE relative to unmodified reference genomes. Nevertheless, RelocaTE produces more correct predictions within 100bp of the true insertion site than ngs_te_mapper, the other purely split-read method despite producing fewer exact predictions than ngs_te_mapper. Thus many of the non-exact RelocaTE predictions within 100bp of the true location are likely to be accurately positioned, but simply not have the correct TSD structure (see below). Like ngs_te_mapper, RelocaTE also appears to have a slightly higher true positive rate for positive strand insertions, with the difference in the number of correct predictions on the positive strand being greater in the exact prediction category.

The average total number of non-reference TE predictions for TEMP is nearly one (0.90), confirming results from unmodified reference genome simulations (see row nine in Additional Table 2) that TEMP makes very few false positive non-reference predictions. Moreover, the total number of non-reference TE predictions for TEMP is the same as the average number that are accurate within 100bp of the true insertion site. These results suggest TEMP is correctly predicting most simulated insertions, but not to base pair accuracy (see below). Some positional inaccuracy is expected for TEMP since not all predictions for this method are supported by split-read evidence. For TEMP, there appears to be no difference in detection ability for TE insertions on the positive or negative strand.

RetroSeq predicted nearly as high an average number of non-reference TE predictions per sample as TEMP, but the proportion predicted correctly was lower than TEMP for all length thresholds. The fact that not all RetroSeq predictions are within 500bp of the true insertion suggests that RetroSeq can produce some false positive predictions of non-reference TE insertions when the sample is not identical to the reference genome, unlike what was observed for simulations of unmodified reference genomes (see row nine of Additional Table 2). Because RetroSeq does not use split-read information, no predictions from this method were exact, however most predictions were generally within 100bp of the true location. For RetroSeq there is a only slight reduction in ability to detect non-reference TE insertions on the negative strand compared with the positive strand at all length thresholds.

PoPoolationTE produces an average of slightly more than one non-reference TE prediction per sample, but this method shows the lowest proportion of true positive predictions at all length thresholds (0.16), suggesting most predictions are false positives. This result supports those obtained from unmodified reference genomes that PoPoolationTE makes approximately one false positive prediction per genome in the absence of any synthetic non-reference TE insertions (see row nine of Additional Table 2). Because PoPoolationTE does not use split-read information and the span predicted by this method is often large (see Additional Figure 1), no predictions made by PoPoolationTE were exact. For PoPoolationTE there appears to be no difference in ability to detect non-reference TE insertions correctly on the positive or negative strand.

TE-locate produced an average of nearly one non-reference TE prediction per sample. However, these include some false positive predictions or at least predictions that are more than 500bp from the actual insertion location. The proportion of correct non-reference TE insertion predicted by TE-locate drops steadily from 500bp to 100bp, with TE-locate predicting the lowest number of correct insertions for any method at the 100bp scale. As with the other read-pair methods, no predictions could be considered exact because TE-locate does not predict a TSD. These numbers indicate that, though the ability of TE-locate to detect the presence of a TE in the general vicinity of its true location is good, the annotation will not be as positionally accurate as other read-pair methods like TEMP or RetroSeq. For TE-locate there appears to be a reduction in detection ability at all thresholds for TE insertions on the negative strand compared with the positive strand.

#### Positional accuracy of non-reference TE predictions

To visualize more clearly the positional accuracy of McClintock component methods, the positions of predicted non-reference insertions were plotted around the known location of synthetic insertions (Figures 2 and 3). Plots were produced for each TE family and method to determine if the family of the synthetic insertion affected results for a particular method. Table 4 showed that for split-read methods, there was no increase in the accuracy at thresholds above 100bp and many predictions were exactly correct. For read-pair methods, it appeared predictions could be several hundred base-pairs from the correct location. As such, split-read (Figure 2) and read-pair (Figure 3) results were plotted on different spatial scales. Since TEMP could use both split-read and read-pair evidence, results for this method were partitioned into two categories for visualization. For a small number of cases, RelocaTE (one location) and PoPoolationTE (five locations) predicted non-reference TE insertions at the same genomic location in multiple samples. These predictions must include false positives based on the fact that each synthetic genome had only a single insertions at different genomic locations. Inclusion of these high-frequency false positive predictions dominated the visualization of results for these two methods, and thus predictions for these six cases were filtered prior generating Figures 2 and 3 (see Methods for details).

**Figure 2.**
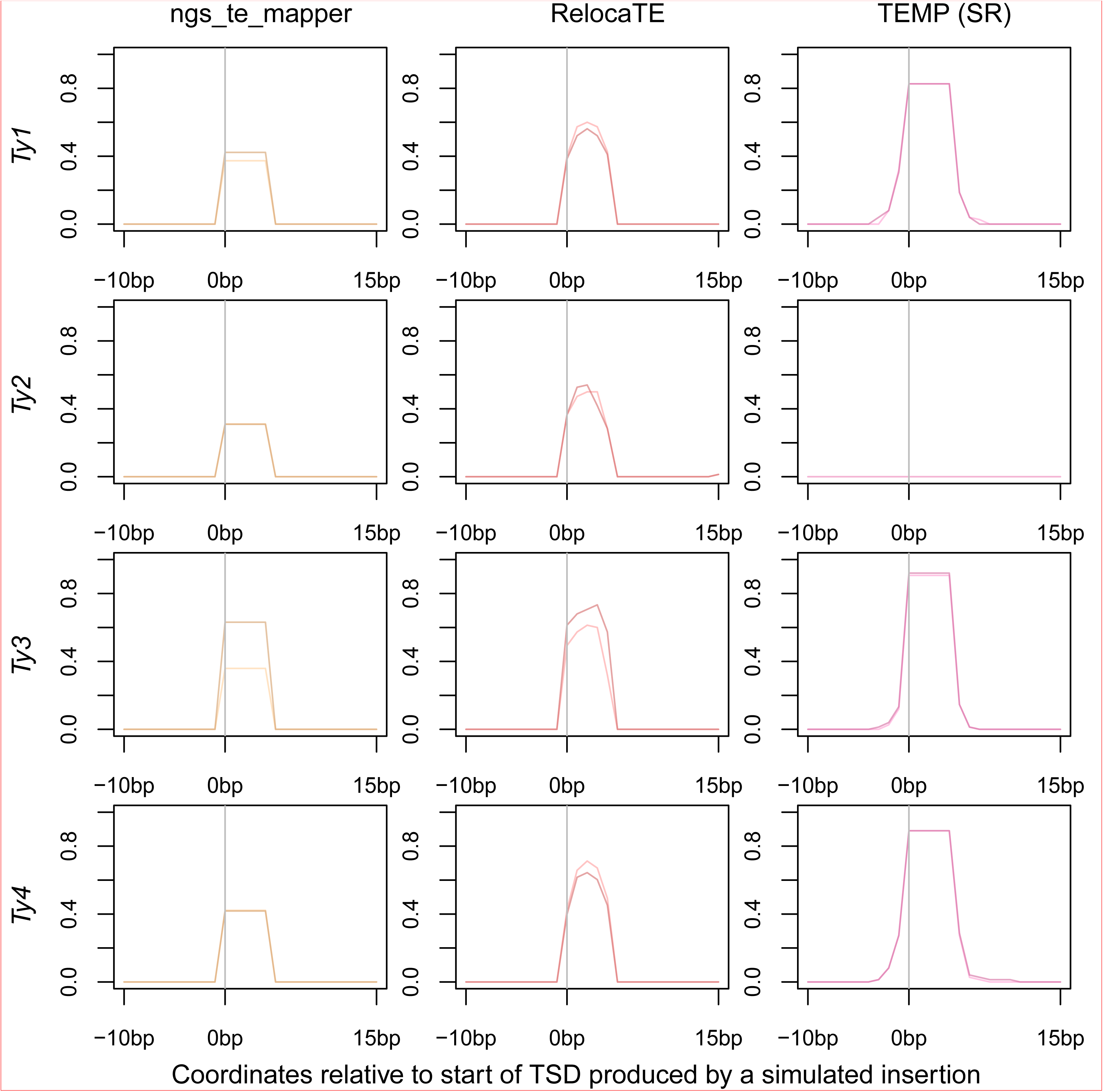
Positional accuracy of non-reference TE insertions made by methods using split-read evidence on single insertion synthetic genomes. Data for TEMP are for predictions that do have split-read evidence and may or may not have read-pair evidence. The location of the synthetic TSD is from position zero to five bp on each plot. The darker line for each method indicates predictions averaged across the 299 simulated genomes with insertions on the positive strand; the lighter line indicates predictions averaged across the 299 simulated genomes with insertions on the negative strand. A value of one would indicate a perfect prediction in all samples since there is one synthetic insertion per genome.

**Figure 3.**
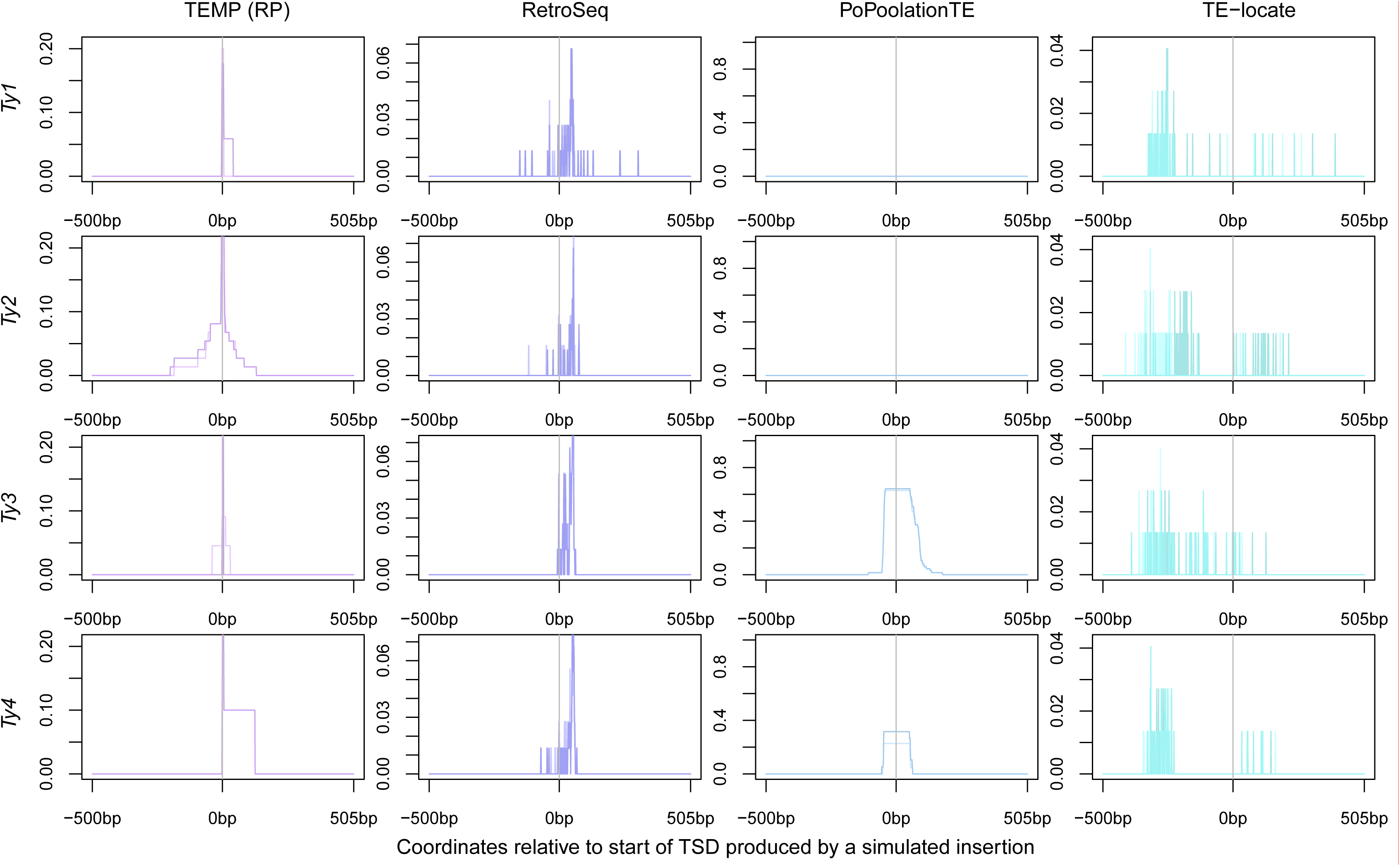
Positional accuracy of non-reference TE insertions made by methods using read-pair evidence on single insertion synthetic genomes. Data for TEMP are for predictions that do not have split-read evidence but do have read-pair evidence. Note that the Y-axes of plots are scaled differently for each method. The location of the synthetic TSD is from position zero to five bp on each plot. The darker line for each method indicates predictions averaged across the 299 simulated genomes with insertions on the positive strand; the lighter line indicates predictions averaged across the 299 simulated genomes with insertions on the negative strand. A value of one would indicate a perfect prediction in all samples since there is one synthetic insertion per genome.

Figure 2 shows that when ngs_te_mapper makes a prediction, it produces the TSD at the correct location, apparently with no TSDs called too long or too short. Direct analysis of TSD length distributions supports this conclusion: for simulated data, ngs_te_mapper always predicts the correct TSD length for non-reference insertions (Additional Figure 2). For *Ty1*, *Ty2* and *Ty4*, ngs_te_mapper detected insertions on the positive or negative strand with similar accuracy. Thus, the main difference in detection rates on the positive and negative strands for ngs_te_mapper observed in Table 4 appears to be for *Ty3* insertions, where many fewer insertions were detected correctly on the negative strand. Because *Ty3* targets an exact location upstream of tRNA genes and the synthetic TE insertions were also made at this location, some synthetic *Ty3* insertions could have been nested in reference copies of *Ty3*, potentially explaining the *Ty3*-specific loss in accuracy. Plots for RelocaTE show that for *Ty1*, *Ty3* and *Ty4*, the predicted TSD is in approximately the correct location but with a coordinate range that is frequently too short (see also Additional Figure 2). As with ngs_te_mapper, RelocaTE shows the biggest difference in ability to detect *Ty3* insertions on the negative strand relative to the positive strand. TEMP split-read predictions for *Ty1*, *Ty3* and *Ty4* are often predicted correctly but with the TSD often annotated to be longer than its true length (see also Additional Figure 2). Surprisingly, TEMP made no predictions for non-reference *Ty2* insertions using split-read evidence, perhaps because of the ambiguous signal arising from the similarity of *Ty1* and *Ty2* LTR sequences. For TEMP, there is no difference in detection ability for insertions on the positive or negative strand for any family.

Results of the positional accuracy for read-pair methods are shown in Figure 3. For *Ty1*, *Ty3* and *Ty4* there were very few insertions (only three per family) that TEMP did not have split-read supporting evidence for, and thus few insertions for these families are plotted in Figure 3. In contrast, all *Ty2* predictions made by TEMP in the single insertion simulations had read-pair evidence. For all families, when only read-pair evidence is used, TEMP generally predicts an insertion at the correct site, but with some slight inaccuracy on either side. The majority of RetroSeq predictions appear to be clustered close to the true insertion locations, but there appears to be a slight bias for RetroSeq to predict insertions 3’ of where the true TE is located on reference genome coordinates. This bias is potentially introduced by the breakpoint determination step of RetroSeq, which always scans in the 5’ to 3’ direction (see section Additional File 1 Description of McClintock Component Methods). PoPoolationTE produced the highest number false positive predictions (Table 4). When these false positive non-reference predictions are filtered from the results, all predictions for *Ty1* and *Ty2* in the windows around simulated insertions are eliminated. The effect of removing false positives is probably most pronounced for Ty1 because Ty1 is the most common TE family in *S. cerevisiae*, and thus would be the most likely family to have a reference insertion with sequence similarity to the synthetic insertion in the vicinity of tRNA genes. PoPoolationTE makes no predictions for Ty2, even including false positives. For Ty3 and Ty4, PoPoolationTE has the capability of producing relatively accurate predictions, albeit with low resolution (nearly 100bp around the true insertion site). For TE-locate, many predictions are made within 500bp of the true insertion, but they are clearly spread further from the true insertion location than other methods. TE-locate also appears to have a slight bias to predict insertions 5’ of the true insertion location on reference genome coordinates.

#### Overlap between methods

To understand the concordance of predictions made by the McClintock components, we investigated the overlap among methods for predictions that were made correctly at the sites of synthetic insertions. As shown in Figures 2 and 3, different methods have different positional accuracy, and thus we used different windows to classify if a method made a “correct” prediction for a known insertion or not. Predictions for ngs_te_mapper, RelocaTE, TEMP (both split-read and read-pair) and PoPoolationTE were classified as correct if the had any overlap with the true location of the TSD, while predictions for RetroSeq and TE-locate were classified as correct if they occurred within a 100 or 500bp window, respectively, of the correct location of the TSD. Neither the orientation nor the TE family was taken into account when classifying a prediction as correct or not. The overlap of correctly detected insertions are shown in Figures 4A and 4B for split-read and read-pair insertions, respectively. The overlap of correct predictions made by all split-read methods versus all read-pair methods is shown in Figure 4C.

**Figure 4.**
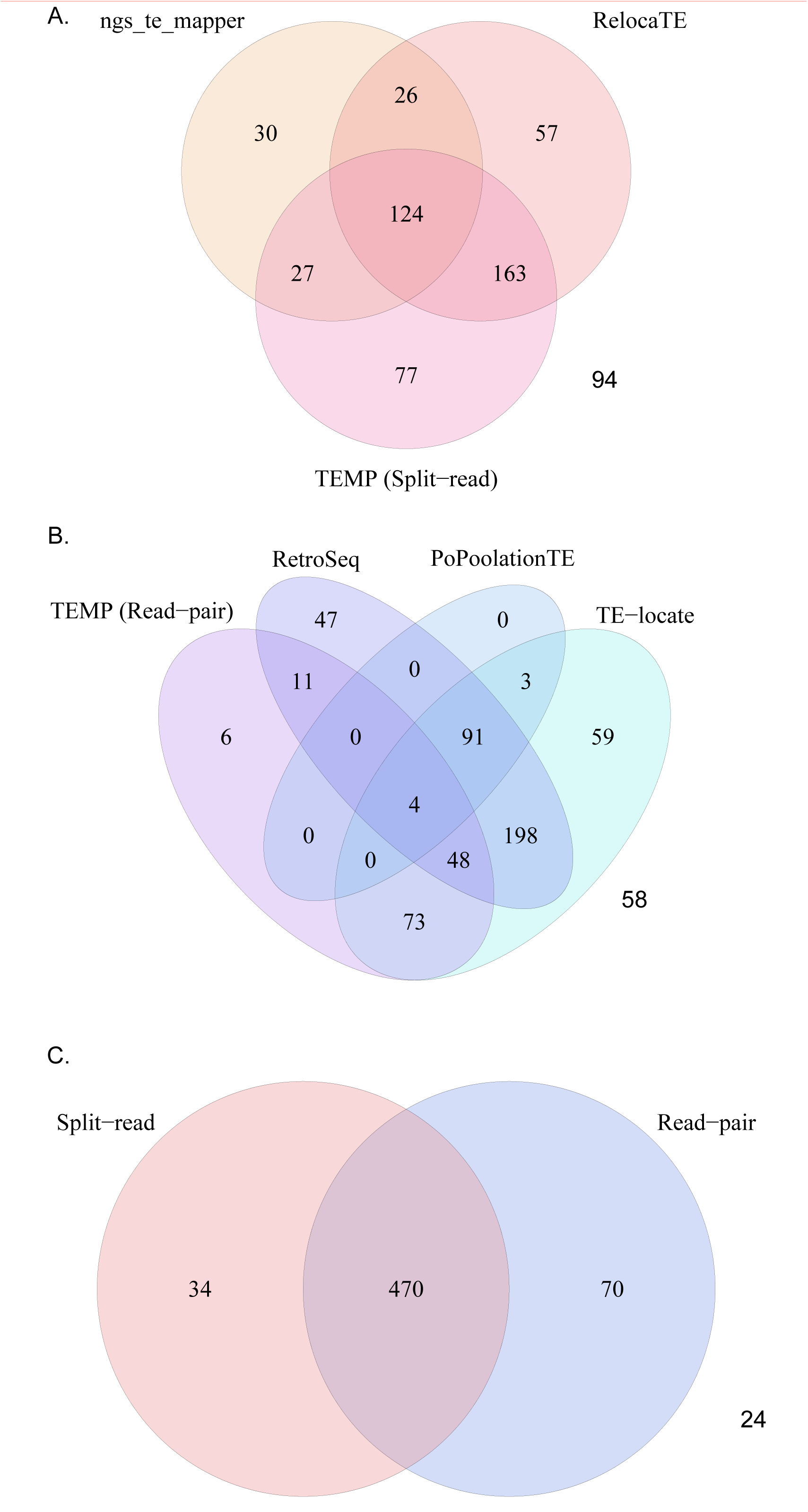
Concordance of correctly predicted non-reference insertions among McClintock component methods. (A) shows the concordance of non-reference predictions by methods that use split-read evidence that overlap with the true location of a synthetic insertion. (B) shows the concordance of non-reference predictions made by methods that use read-pair evidence that either overlap (TEMP, PoPoolationTE), or are within 100bp (RetroSeq), or 500bp (TE-locate), of the the true location of a synthetic insertion. (C) shows the concordance of correctly predicted synthetic non-reference TEs with split-read or read-pair evidence. Predictions for TEMP were partitioned based on whether they had split-read evidence (split-read) or not (read-pair). Counts in all diagrams total 598, the number of simulated samples with single synthetic insertions.

Figure 4A shows that the majority of split-read predictions are supported by at least two methods (n=340, 57%) but that each method made many correct TE predictions that were not made by any other method. RelocaTE and TEMP made a greater number of correct overlapping predictions with each other than either of these method did with ngs_te_mapper. Figure 4A also shows that 16% (n=94) of synthetic insertions were not predicted by any split-read method at the threshold of positional accuracy used here. Figure 4B shows that the vast majority of synthetic TE insertions (n=428, 72%) are predicted by at least two of the read-pair methods, but that only 24% (n=143) of insertions are supported by three or more methods. RetroSeq and TE-locate make the highest number of unique correct predictions. Approximately 10% (n=58) of synthetic insertion samples were not predicted by any read-pair method at the threshold of positional accuracy used here. Finally, Figure 4C shows that, while the overwhelming the majority are predicted by at least one split-read and one read-pair method (n=470, 79%), there are many insertions that are only predicted using one type of evidence or the other (n=104, 17%) given the thresholds of positional accuracy used here. Nevertheless, use of all six methods recovers nearly 96% of synthetic insertions, demonstration the utility of integrating multiple TE identification methods enabled by McClintock.

### Application of McClintock to 93 yeast genomes

The previous sections presented results on the accuracy of McClintock component methods on simulated resequencing data. Simulations are useful for testing methods under controlled settings, but do not capture all aspects of how methods perform when applied to real data. Since much is known about the expected insertion preferences of TEs in *S. cerevisiae* [44, 46–55], analysis of real WGS datasets can be used as an alternative approach to evaluate if McClintock component methods can recapitulate the known genome biology of yeast TEs. To do this, we analyzed 93 high-coverage *S. cerevisiae* WGS datasets from Strope *et al.* [42] using McClintock to generate TE predictions for all six component methods. Figure 3 and Additional Figure 5 show how many of non-reference and reference TEs per strain, respectively, are detected by the different McClintock component methods across all 93 samples. In general, split-read methods predict between 5-20 non-reference TE insertions per strain, whereas read-pair methods predict approximately 40-100 non-reference TE insertions per strain (Figure 3). Numbers of reference TEs predicted per strain in real data (Additional Figure 5) are generally lower than in simulated genomes (Table 4 and Additional Table 1). The exceptions to this pattern are TEMP and PoPoolationTE, which show similar or higher numbers of reference TE predictions per strain in real data relative to simulations. We note that for a few strains in the Strope *et al.* [42] dataset, TE-locate predicted several hundred non-reference insertions; these strains did not appear to be outliers in terms of their non-reference TE content based on other methods (results not shown).

**Figure 5.**
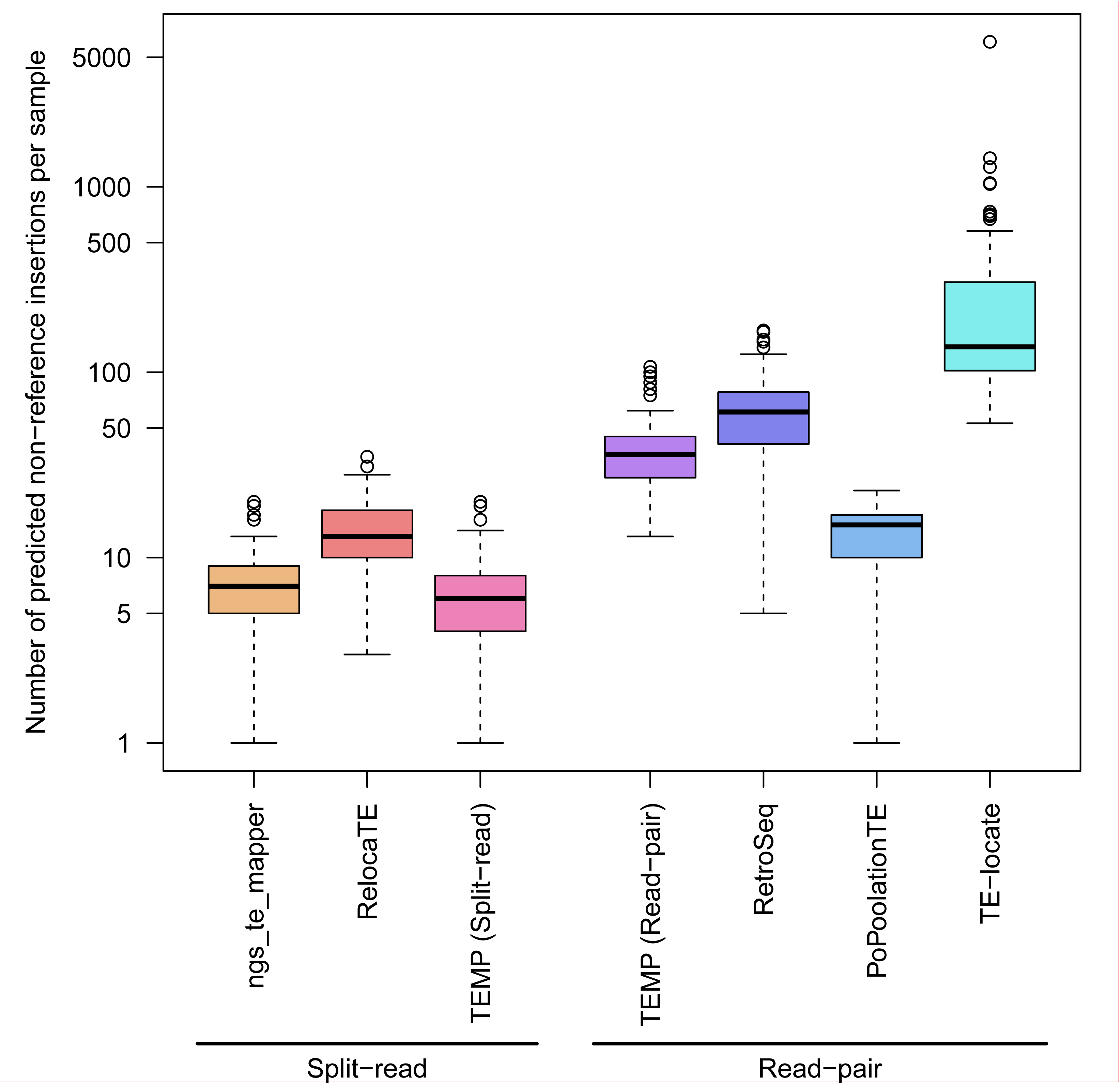
Numbers of non-reference TE insertions per strain predicted by McClintock component methods in real yeast genomes. Predictions for TEMP were partitioned based on whether they had split-read evidence (split-read) or not (read-pair). Data are from 93 yeast strains taken from Strope *et al.* [42]. Methods are classified based on whether they use split-read or read-pair evidence to make a non-reference TE prediction. The box plot is shown on a log_10_ scale. The thick line indicates the median, the colored box is the interquartile range, the whiskers mark the most extreme data point which is no more than 1.5 times the interquartile range from the box, and the circles are outliers. Note that for TE-locate, several outlier samples generated hundreds of predicted non-reference TE insertions.

We evaluated the quality of non-reference TE predictions made by McClintock component methods on the Strope *et al.* [42] dataset using three aspects of the known biology of TEs in *S. cerevisiae*: (i) activity of families, (ii) tRNA targeting, and (iii) TSD length. Our expectations based on prior knowledge of yeast TE biology are that methods that make high quality non-reference TE predictions should (i) show few non-reference predictions for inactive TE families (*Ty3_1p* and *Ty5*), (ii) show a high proportion of non-reference predictions in the vicinity of tRNA genes, and (iii) show characteristic 5 bp TSDs for non-reference predictions made by split-read methods.

#### Prediction of active and inactive families

Table 5 shows numbers of non-reference TE predictions made by McClintock component methods across all strains in the Strope *et al.* [42] dataset. As expected, all methods predicted multiple non-reference insertions for TE families that are known to be active in this species. Additionally, ngs_te_mapper and TEMP make no non-reference TE predictions for both inactive families in *S. cerevisiae*, supporting simulation results above that show these methods have low false positive rates. RelocaTE makes non-reference TE predictions for *Ty3_1p* but not *Ty5*, PoPoolationTE makes non-reference TE predictions for *Ty5* but not *Ty3_1p*, and both RetroSeq and TE-locate predict non-reference insertions for *Ty3_1p* and *Ty5*. RelocaTE is the only split-read method that predicts non-reference insertions for an inactive family, suggesting that split-read methods generally have higher ability to discriminate active from inactive TE families. Compared to the total numbers predicted for other active TE families, the three pure read-pair methods predicted fewer non-reference insertions for both *Ty3_1p* and *Ty5*, suggesting false positive are not so numerous as to overwhelm true signal. The one exception is for TE-locate, which predicted relatively high numbers of *Ty5* insertions, which is likely related to the outlier samples noted above where TE-locate predicts hundreds of presumably false-positive insertions.

**Table 5:** Number and location of non-reference TEs predicted by McClintock component methods in 93 yeast genomes from Strope *et al.* [42]. Each cell shows the number of non-reference TEs predicted in tRNA regions followed by the total number of non-reference TEs predicted genome wide. Data are for numbers of insertions, not numbers of non-redundant insertion sites, so TE insertion alleles present in more than one sample are counted independently. A prediction is counted in a tRNA region if any portion of the annotation is within 1000bp upstream and 500bp downstream of the tRNA start site, taking into account the orientation of the tRNA gene. The first column applies the same analysis to the reference TE annotations from Carr *et al.* [45]. N. A. indicates that no non-reference TE insertions were predicted by a method for that TE family.

|  | **Carr** | **ngs_te_mapper** | **RelocaTE** | **TEMP** | **RetroSeq** | **PoPoolationTE** | **TE-locate** |
| --- | --- | --- | --- | --- | --- | --- | --- |
| *Ty1* | 218/313 (70%) | 93/101 (92%) | 15/18 (83%) | 827/1093 (76%) | 1854/2835 (65%) | 139/194 (72%) | 2082/16388 (13%) |
| *Ty2* | 30/46 (65%) | 58/77 (75%) | 303/425 (71%) | 1343/1853 (72%) | 839/1169 (72%) | 27/36 (75%) | 1110/8132 (14%) |
| *Ty3* | 43/45 (96%) | 378/387 (98%) | 670/678 (99%) | 991/1008 (98%) | 1299/1445 (90%) | 1006/1013 (99%) | 1748/3813 (46%) |
| *Ty3_1p* | 12/15 (80%) | 0/0 (N.A.) | 23/23 (100%) | 0/0 (N.A.) | 12/16 (75%) | 0/0 (N.A.) | 83/86 (97%) |
| *Ty4* | 29/49 (59%) | 95/118 (81%) | 143/190 (75%) | 259/310 (84%) | 238/292 (82%) | 15/20 (75%) | 324/1083 (30%) |
| *Ty5* | 0/15 (0%) | 0/0 (N.A.) | 0/0 (N.A.) | 0/0 (N.A.) | 3/74 (4%) | 0/12 (0%) | 0/887 (0%) |

#### Predicted insertions in tRNA regions

Active TE families in *S. cerevisiae* are known to target tRNA genes [44, 47–50, 53–55]. The highest density of *Ty1* and *Ty2* insertions are in the 200bp upstream of the tRNA transcription start site [44, 47, 50, 53, 54]. *Ty3* targets a specific location just upstream of tRNA gene transcription start sites [44, 48, 49, 55]. Patterns of *Ty4* insertions have not been experimentally determined, although the locations of insertions in the reference genome suggest a similar pattern to that of *Ty1* and *Ty2* [44].

To evaluate if non-reference TE insertions predicted by McClintock component methods show expected hallmarks of tRNA targeting, we plotted locations of non-reference TE insertions identified in the Strope *et al.* [42] strains using split-read evidence and read-pair evidence in Figures 6 and 7, respectively. The expected profiles of insertion into tRNA gene regions is observed for all Ty families for ngs_te_mapper, RelocaTE, TEMP and RetroSeq, albeit with the different levels of resolution that are characteristic of each method. Consistent with simulation data (Figure 2), TEMP appears to have difficulty predicting *Ty2* using split-read data in real yeast genomes, and this effect also appears to impact prediction of *Ty1* insertions using split-read data in real data (Figure 6). PoPoolationTE can predict meaningful profiles of insertion for *Ty3* and *Ty4* (Figure 7), as expected based on simulation data (Figure 3). However, in contrast to simulation data where only putative false positives are predicted (Figure 3), PoPoolationTE also predicts non-reference insertions for *Ty1* and *Ty2* in real data (Figure 7). Since PoPoolationTE predicts reference and non-reference insertions in the same way, and since many *Ty1* and *Ty2* insertions exist in the reference genome upstream regions of tRNA genes, it is possible that these *Ty1* and *Ty2* insertions predicted in the Strope *et al.* [42] dataset are actually reference insertions that are mislabelled by PoPoolationTE as non-reference insertions. Finally, non-reference insertions predicted by TE-locate are only weakly enriched in tRNA regions for all families, and the positional profiles produced by TE-locate are shifted relative to expectations and predictions made by other methods.

**Figure 6.**
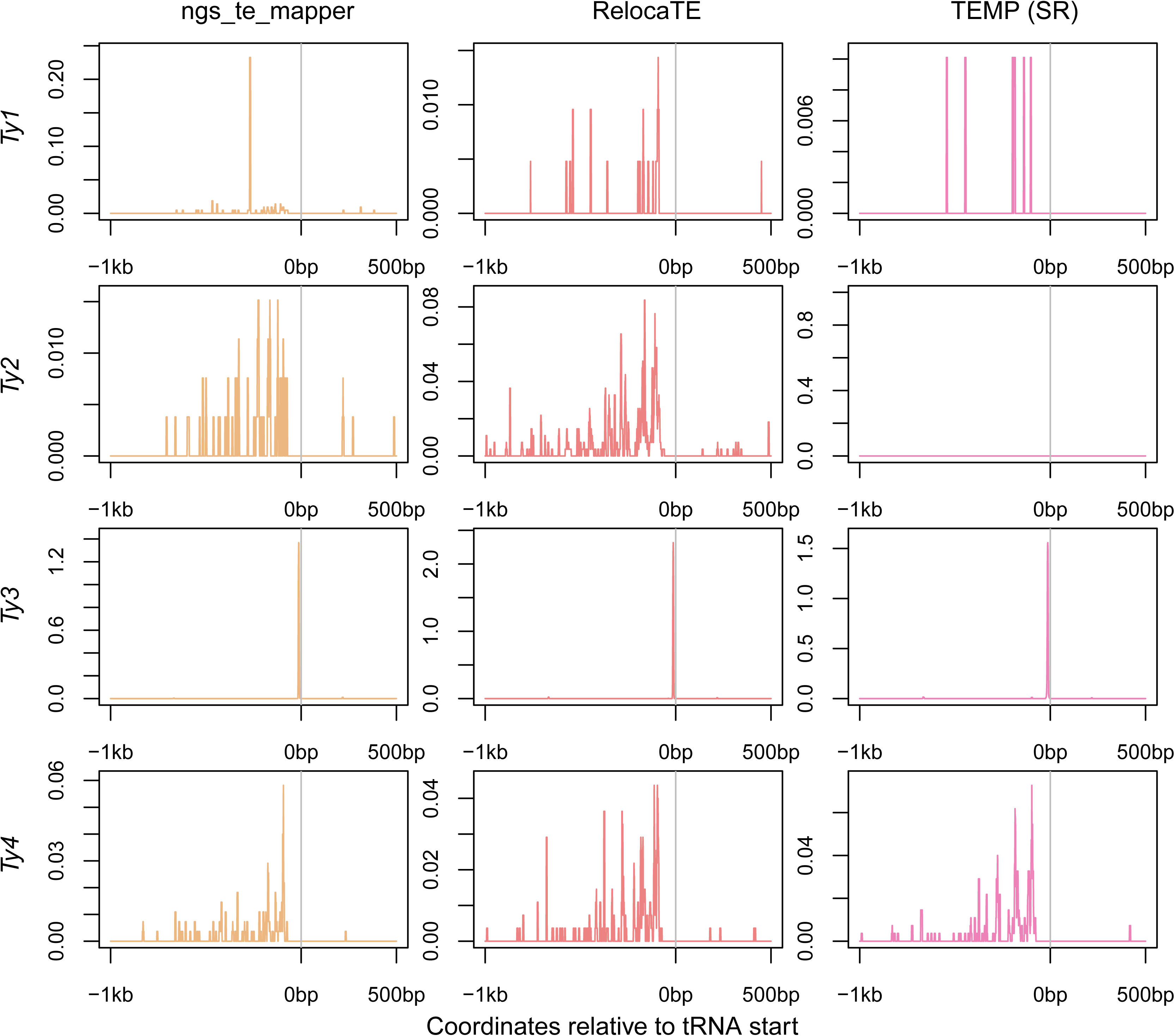
Locations of non-reference TE predictions relative to tRNA genes made by methods using split-read evidence on real yeast genomes. Data for TEMP are for predictions that do have split-read evidence and may or may not have read-pair evidence. The transcription start site of each tRNA gene is aligned at position zero, taking into tRNA gene account orientation, for a window extending 1kb upstream and 500bp downstream. The frequency of a prediction at each base is counted across all 93 strains in the Strope *et al.* [42] dataset, then averaged across the 299 tRNA genes and plotted as a line for each method and TE family. These plots show all predictions and therefore include allelic predictions present in more than one strain. Also, any given strain may have more than one insertion at the same relative location in a tRNA gene, and thus the scale for these plots can go above one.

**Figure 7.**
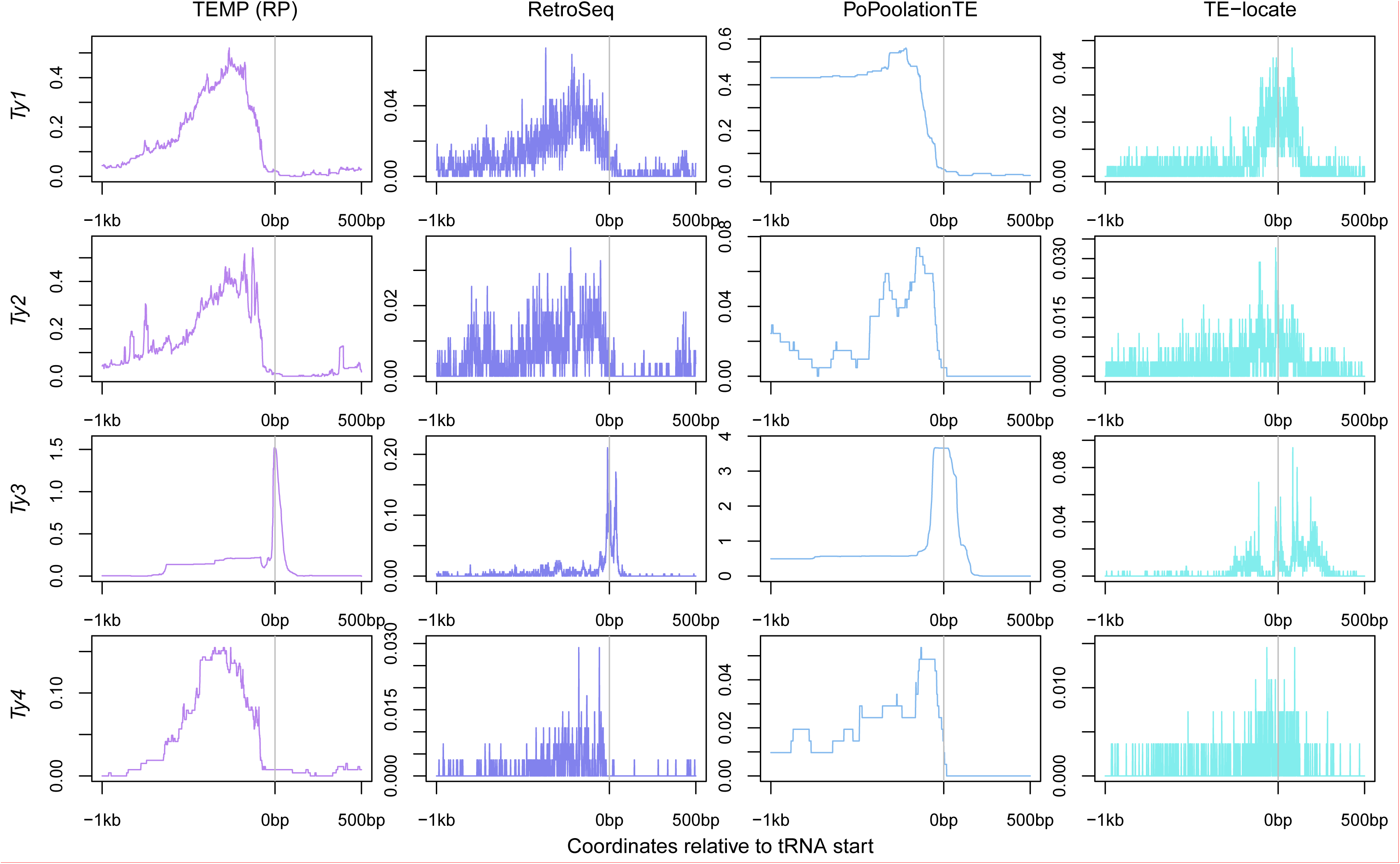
Locations of non-reference TE predictions relative to tRNA genes made by methods using read-pair evidence on real yeast genomes. Data for TEMP are for predictions that do not have split-read evidence but do have read-pair evidence. The transcription start site of each tRNA gene is aligned at zero on the plots, taking into account orientation, for a window extending 1kb upstream and 500bp downstream. The frequency of a prediction at each base is counted across all 93 strains in the Strope *et al.* [42] dataset, then averaged across the 299 tRNA genes and plotted as a line for each method and TE family. These plots show all predictions and therefore include allelic predictions present in more than one strain. Also, any given strain may have more than one insertion at the same relative location in a tRNA gene, and thus the scale for these plots can go above one.

To quantify the proportion of non-reference TEs that were predicted in tRNA regions, we counted predictions 1000bp upstream and 500bp downstream of a tRNA gene, taking into account the orientation of the tRNA gene but not the orientation of the TE insertion. The expected percentage of TEs located in these regions if they were inserted randomly in the genome would be 0.037% ((299 tRNA genes × 1500bp window) ÷ 12,162,995bp genome). Previous analyses of tRNA targetting of TEs in the *S. cerevisiae* reference genome [44] assessed whether TEs were within 750bp of a tRNA gene or other RNA polymerase III gene (excluding other intervening TE sequences). Here we use extended regions for tRNA targeting based on the inaccuracy in non-reference predictions observed for some methods in the simulations above. For comparison with previous results, we first applied our definition of tRNA targeting to the reference TE annotation from Carr *et al.* [45] (Table 5). Estimated proportions of *Ty* elements in tRNA regions for the Carr *et al.* [45] reference annotation are lower than those reported by Kim *et al.* [44], however, they still show highly biased targeting towards tRNA regions.

Non-reference TE predictions of all four active *Ty* elements show the expected enrichment in tRNA regions for each McClintock component method (Table 5). For all methods, *Ty3* is the active TE family most strongly associated with tRNA regions, consistent with experimental data and observations based on the reference genome [44, 48, 49, 55]. Split-read methods predict a higher proportion of non-reference TEs in tRNA regions relative to expectations based on TEs in the reference genome. For read-pair methods, at least one TE family showed a lower proportion of non-reference TEs in tRNA regions relative to reference TEs. We interpret this observation to be due to the lower positional accuracy of read-pair methods. TE-locate consistently predicted the lowest number of TEs in tRNA regions for active *Ty* families, though predictions for this method still showed a bias towards insertion in tRNA regions relative to random insertions. We interpret the low tRNA enrichment for TE-locate to be a consequence of the low positional accuracy of read pair methods combined with the presence of outlier samples for this method that have very high numbers of non-reference predictions.

As discussed above, non-reference predictions were made by RelocaTE, RetroSeq, and TE-locate for the inactive *Ty3_1p* family. Despite most likely being false positives, these predictions were predominantly in tRNA regions, suggesting they could either be non-reference *Ty3* insertions that are miscalled as non-reference *Ty3_1p*, or reference *Ty3_1p* insertions called as non-reference *Ty3_1p* insertions. Non-reference predictions were also made by RetroSeq, PoPoolationTE and TE-locate for the inactive *Ty5* family. The majority of these predictions are made outside of the tRNA regions, as is expected based on the known location of *Ty5* insertions in the reference genome prior knowledge about *Ty5* target preferences [44, 52, 56]. These non-reference TE predictions may be false positives (possibly caused by mapping inconsistencies in heterochromatic regions where *Ty5* elements typically insert) or real non-reference “insertions” present in sequenced strains that arose by recombination events rather than transposition events [57].

#### Prediction of TSDs by split-read methods

Finally, we evaluated the performance of split-read methods to predict the known TSD lengths of active yeast *Ty* families in real WGS data. All available experimental and genomic indicates that active yeast *Ty* families create five bp TSDs on insertion [44, 46, 48, 51, 52]. TSD length distributions of unique insertion sites are shown for *Ty1*, *Ty2*, *Ty3*, and *Ty4* in Additional Figure 4. As observed in simulated data (Figure 2), ngs_te_mapper predictions had the highest proportion of correct TSD lengths predicted per family. However, in contrast to simulated data, ngs_te_mapper can infrequently make incorrect TSD length predictions in real data. Confirming simulation results, RelocaTE generally under-predicts the length of TSDs, and TEMP consistently over-predicts the lengths of TSDs for all families in real data. For all split-read methods, the modal value of the TSD length distribution reflects the true TSD length for all families. Thus, the modal TSD length provided by each of the split-read methods yields biologically meaningful inferences about TSD structure.

## Conclusions

Here we described McClintock, an integrated pipeline for detection TE insertions in WGS resequencing data. McClintock offers many advantages relative to running multiple TE detection methods in isolation. Specific versions of compatible software dependencies required to run each method are fully documented, allowing users to easily set up their environment. The number of input files required to run all methods is reduced and complex processing of input files to create the correct custom formats and file relationships is automated. In addition, the pipeline is structured to allow parallel computations for multiple samples, so population datasets can be analyzed more quickly. Finally, results from individual methods are standardized to facilitate comparisons across methods and easy visualization in the UCSC genome browser. Overall, McClintock greatly lowers the barriers to to running multiple TE detection methods, allowing users to gain more insight into how various methods work for their samples. McClintock does not currently include all published TE detection methods, although additional methods can be easily incorporated into the pipeline due to the flexible architecture and open source nature of the project.

In addition, we have applied McClintock to simulated and real yeast WGS samples to evaluate the performance of McClintock component methods. Simulations on the unmodified *S. cerevisiae* reference genomes reveal that sequencing coverage influences detection of reference TEs, but that recovery of reference TE insertions and false positive rates for non-reference TE insertions are generally low even at high sequencing coverage. Simulations on *S. cerevisiae* reference genomes including a single non-reference insertion showed that pure split-read methods may detect fewer TE insertions than read-pair methods, but they have much higher positional accuracy. Single insertion simulations also revealed that the TE family affects the ability of methods to detect non-reference TE insertions. We find substantial difference in the ability of McClintock component methods to detect subsets of non-reference insertions in the yeast genome, but that by combining multiple methods that use split-read and read-pair data, non-reference TEs at nearly all biologically-realistic locations can be detected in simulated data. Finally, application of McClintock to a large sample of real yeast genomes reveals that most but not all McClintock component methods can recovery known aspects of TE biology in yeast such as family activity status, tRNA gene targeting, and TSD structure. Together, our results suggest that even in the context of a simplified model eukaryotic genome like *S. cerevisiae*, current TE detection methods using short-read data do not provide comprehensive recovery of all TE insertions in WGS resequencing samples. Further performance studies in other genomic contexts including newer methods not currently included in McClintock are needed to generalize the results presented here, and to provide a road map for developing more advanced systems for the detection of TEs in unassembled short-read genomic data.

## Methods

### Analysis of simulated WGS datasets with single artificial TE insertions

To investigate the performance of McClintock component methods on data containing known, non-reference TE insertions, we created simulated *S. cerevisiae* genome, each containing one synthetic non-reference TE insertion in an otherwise unmodified *S. cerevisiae* reference genome. Since active *S. cerevisiae* TEs (*Ty1*, *Ty2*, *Ty3* and *Ty4*) are known to target tRNA genes [44, 47–50, 53–55], each synthetic insertion was placed upstream of a different annotated tRNA gene in the reference genome, taking the orientation of the tRNA gene into consideration. The annotation for 299 tRNAs was extracted from the SGD genome annotation for sacCer2 (SGD version R61.1.1). *Ty1*, *Ty2* and *Ty4* have been shown to insert predominantly within the first 200bp upstream of tRNA genes, and *Ty3* appears to target more specifically the region of RNA polymerase III transcription initiation, 16 or 17 nucleotides from the 5’ ends of tRNA genes [44, 47-50, 53-55]. All active *S. cerevisiae* TEs produce five bp TSDs on insertion [44, 46, 48, 51, 52]. To mimic these insertion preferences in our simulations, *Ty1*, *Ty2*, *Ty3* and *Ty4* were alternately selected for insertion, a five bp TSD was created (either 200bp to 195bp upstream of a tRNA gene for *Ty1*, *Ty2* and *Ty4*, or 17bp to 12bp upstream of tRNA genes for *Ty3*), and the corresponding full length *Ty* canonical sequence was inserted in the reference genome. 299 insertions were produced with the TE sequence inserted on the positive strand of the genome, and 299 were produced with the TE sequence reverse complemented to test the effects of TE orientation on method performance.

We simulated resequencing of single-insertion synthetic genome using WgSim (https://github.com/lh3/wgsim) [58] with parameters that resemble the properties of Illumina sequencing (as described by [59]). Read lengths were chosen to be 101 bases each with an insert size of 300 bases and 100X coverage to mimic the properties of a large sample of WGS datasets collected by Strope *et al.* [42] used in our analysis of real yeast genomes (see below). To generate an average read depth of 100X across the length of sacCer2 reference genomes with additional single TE insertions, *in silico*’ WGS samples were created with 6,024,220 read pairs for *Ty1* insertions, 6,024,237 read pairs for *Ty2* insertions, 6,023,936 read pairs for *Ty3* insertions, and 6,024,369 read pairs for a *Ty4* insertion.

McClintock (version e945d20da22dc1186b97960b44b86bc21c96ac27) was run on each of these simulated datasets using reference TE annotations and canonical TE sequences from Carr *et al.* [45], plus a manually produced hierarchy file based on the reference TE annotation in [45]. We used the standard, unmodified reference genome sequence option of McClintock for these single synthetic insertion simulations. The mean of the number of non-reference and reference TEs predicted per sample was calculated across all 299 simulated samples for each strand. The proportion of correct predictions of non-reference TEs was calculated at four thresholds of accuracy: (i) requiring the exact TSD to be annotated correctly, (ii) requiring a prediction to be within a 100bp window either side of the TSD, (iii) within a 300bp (the insert size of the simulated sequencing dataset) window either side of the TSD, or (iv) within a 500bp window either side of the TSD. BEDtools window [60] was used to calculate correct predictions within the given windows. A prediction was classified as exactly correct only if the same TE family was predicted to occur at the exact coordinates of the TSD of the synthetic TE insertion location. For non-exact overlaps, BEDtools window allows a permissive definition of a true positive, where a correct TE prediction is counted when any part of a predicted insertion falls within the given threshold distance if the correct TE family is predicted. The orientation of a predicted insertion was not taken into account for determining a correct prediction because some methods do not predict orientation.

To visualize the accuracy of non-reference TE predictions, the results files for the 299 positive strand and 299 negative strand single insertion samples were converted into two BigWig files (one for each strand) using BEDtools and wigToBigWig [61]. This was performed for each TE family and each component method of McClintock. SeqPlots [62] was then used to produce plots of the genome coverage of predictions for each TE family, centered around the simulated insertion locations for that family. Visualization of predicted insertions negative strand simulations were reverse complemented and depicted on the same plot as positive strand simulations in different colors. Plots were centered on the five bp TSD and extended ±10bp for split-read methods, and ±500bp for read-pair methods, respectively. Results for TEMP were partitioned based on whether or not split-read support was available for a prediction. Prior to visualization, we attempted to filter out any obvious false positive predictions using the fact that each synthetic insertion location should only be predicted in one simulated sample. Thus, any locations where a predicted non-reference insertion was observed across multiple simulated samples indicated a potential false positive. This filtering was necessary to prevent a non-reference insertion that was predicted by RelocaTE in the same location in 149 single synthetic insertion samples from dominating the visualization for this component method. False positive filtering prior to visualization only affected five other potential insertions for PoPoolationTE, and thus this filtering procedure does not substantially alter positional accuracy results. To further investigate the accuracy of TSDs predicted by split-read methods, the length of the predicted TSD was plotted for each active yeast TE family. To be consistent with analysis of real yeast genomes (see below) and to mitigate effects of false positive predictions found at the same site in multiple samples, TSD lengths predicted in simulated data were only plotted for unique insertion sites rather than all insertions.

To investigate the concordance of non-reference TE predictions made by different McClintock component methods, we first determined whether or not each method had made a “correct” prediction in each of the simulated samples with a synthetic TE insertion. Predictions for ngs_te_mapper, RelocaTE, TEMP (both split-read and read-pair), and PoPoolationTE were classified as correct if they overlapped with the true location of the TSD. Predictions for RetroSeq and TE-locate were classified as correct if they occurred within a 100 or 500bp window of the correct location of the TSD, respectively. The orientation of a prediction was not taken into account when classifying a prediction as “correct” or not, because not all methods predict strand. The overlap of these correct predictions was then plotted as venn diagrams using jvenn [63], comparing split-read methods, read-pair methods and finally the total set of correct predictions from all split-read versus all read-pair methods.

### Analysis of real WGS datasets

To assess the relative performance characteristics of the component methods on real data, McClintock was run on a large sample of *S. cerevisiae* datasets from Strope *et al.* [42] that includes 93 *S. cerevisiae* strains from different geographical locations and clinical origins. Strope *et al.* [42] samples were sequenced on an Illumina HiSeq 2000 with paired-end reads of 101 bases each, an average insert size of 300 bases, and a median coverage of greater than 117X. We used these general library characteristics in our single synthetic insertion simulations (above), to allow more direct comparison with analysis of these real yeast genomes. The raw fastq files for the 93 sequenced strains were obtained from the EBI Sequence Read Archive (SRA072302).

McClintock (version 354acec977e37c354f6f05046940b0dabf09b331) was run on each of these samples using reference TE annotations and canonical TE sequences from Carr *et al.* [45], and a manually produced hierarchy file based on the annotation in [45]. The McClintock version used for analysis of real yeast data differs slightly from that used for simulated data in terms of three small improvements that were required to handle variation in sample names (for ngs_te_mapper) and differences in read lengths of paired end fragments (for PoPoolationTE) that were encountered when analyzing real yeast genome data. We used the standard, unmodified reference genome sequence option of McClintock for these analyses. The average number of non-reference and reference TEs predicted per strain was plotted as box plots for each method. In addition, the total numbers of non-reference and reference TE insertions per TE family were summarized across all strains for each McClintock component method, both genome-wide and in tRNA gene regions.

To biologically validate results of different component methods of McClintock, we took advantage of the fact that *Ty* elements are known to insert in close proximity to tRNA genes in *S. cerevisiae* [44], 47-50, 53-[55]. A prediction was counted as within a tRNA gene region if any part of the annotation was with 1000bp upstream or 500bp downstream of the transcription start site of one of the 299 annotated tRNA genes, taking tRNA gene orientation into account. To visualize the patterns of non-reference TE predictions around tRNA genes, all results for all 93 samples were converted to a single genome coverage bigWig file for each TE and each component method. SeqPlots [62] was used to produce plots of the genome coverage averaged across the 299 tRNA genes. Plots were centered on the start of the tRNA gene and extended 1000bp upstream and 500bp downstream, taking into account the orientation of each tRNA gene. Results for TEMP were subset into two groups based on whether split-read support for a prediction was available or not.

The lengths of TSDs for non-reference TE insertions predicted by the split-read methods were plotted by TE family. To prevent any non-reference TE insertions present at the same location in multiple samples from biasing the results, only unique insertion sites were plotted. If a method called an insertion at nearly the same location but with a longer or shorter TSD in different samples, these were classed as unique sites.

## Declarations

### Availability of Data and Materials

Supporting datasets of McClintock predictions for real yeast genomes are available as Additional File 2.

### Availability of supporting source code and requirements

Project name: McClintock

Project home page: https://github.com/bergmanlab/mcclintock

Operating system(s): LINUX

Programming language: BASH, PERL

License: FreeBSD

### Abbreviations

LTR: long terminal repeat
TE: transposable element
TSD: target site duplication
WGS: whole-genome shotgun

### Ethics approval and consent to participate

Not applicable.

### Consent for publication

Not applicable.

## Author’s contributions

MGN developed the McClintock pipeline, wrote the simulation and analysis code, performed the analysis, and drafted the manuscript; RSL developed ngs_te_mapper; CMB conceived of and supervised the project, performed the analysis, and drafted the manuscript.

## Additional Files

Additional File 1 contains a combined supplement including: detailed descriptions of McClintock components; details of post-processing of component method output; methods, results and discussion of running McClintock on simulated resequencing data created for unmodified *S. cerevisiae* reference genomes; Additional Tables 1–2; and Additional Figures 1–4. Additional File 2 contains McClintock predictions for real yeast genomes on sacCer2 coordinates in BED format.

## Acknowledgements

We thank Matthew Ronshaugen, Douda Bensasson and members of the Bergman Lab and for helpful comments throughout the project; Robert Kofler, Thomas Keane, Alexander Platzer, So a Robb, Jiali Zhuang for code fixes and insights into McClintock component methods; and Nick Gresham for high-performance cluster computing assistance.

## References

1. Bergman, C.M., Quesneville, H.: Discovering and detecting transposable elements in genome sequences. Brief Bioinformatics 8(6), 382–92 (2007). doi:10.1093/bib/bbm048

2. Saha, S., Bridges, S., Magbanua, Z.V., Peterson, D.G.: Computational Approaches and Tools Used in Identification of Dispersed Repetitive DNA Sequences. Tropical Plant Biol. 1(1), 85–96 (2008). doi: 10.1007/s12042-007-9007-5. Accessed 2016-12-14.

3. Lerat, E.: Identifying repeats and transposable elements in sequenced genomes: how to find your way through the dense forest of programs. Heredity (Edinb) 104(6), 520–533 (2010). doi: 10.1038/hdy.2009.165

4. Hoen, D.R., Hickey, G., Bourque, G., Casacuberta, J., Cordaux, R., Feschotte, C., Fiston-Lavier, A.-S., Hua-Van, A., Hubley, R., Kapusta, A., Lerat, E., Maumus, F., Pollock, D.D., Quesneville, H., Smit, A., Wheeler, T.J., Bureau, T.E., Blanchette, M.: A call for benchmarking transposable element annotation methods. Mobile DNA 6, 13 (2015). doi: 10.1186/s13100-015-0044-6. Accessed 2016-01-28.

5. Ewing, A.D.: Transposable element detection from whole genome sequence data. Mobile DNA 6, 24 (2015). doi:10.1186/s13100-015-0055-3. Accessed 2016-01-28.

6. Sackton, T.B., Kulathinal, R.J., Bergman, C.M., Quinlan, A.R., Dopman, E.B., Carneiro, M., Marth, G.T., Hartl, D.L., Clark, A.G.: Population genomic inferences from sparse high-throughput sequencing of two populations of Drosophila melanogaster. Genome Biol Evol 1, 449–65 (2009). doi: 10.1093/gbe/evp048

7. Ewing, A.D., Kazazian, H.H.: High-throughput sequencing reveals extensive variation in human-specific L1 content in individual human genomes. Genome Research 20(9), 1262–70 (2010). doi: 10.1101/gr.106419.110

8. Ewing, A.D., Kazazian, H.H.: Whole-genome resequencing allows detection of many rare LINE-1 insertion alleles in humans. Genome Research (2011). doi: 10.1101/gr.114777.110

9. Hormozdiari, F., Hajirasouliha, I., Dao, P., Hach, F., Yorukoglu, D., Alkan, C., Eichler, E.E., Sahinalp, S.C.: Next-generation VariationHunter: combinatorial algorithms for transposon insertion discovery. Bioinformatics 26(12), 350–7 (2010). doi: 10.1093/bioinformatics/btq216

10. Quinlan, A.R., Clark, R.A., Sokolova, S., Leibowitz, M.L., Zhang, Y., Hurles, M.E., Mell, J.C., Hall, I.M.: Genome-wide mapping and assembly of structural variant breakpoints in the mouse genome. Genome research 20(5), 623–35 (2010). doi: 10.1101/gr.102970.109

11. Fiston-Lavier, A.-S., Carrigan, M., Petrov, D.A., González, J.: T-lex: a program for fast and accurate assessment of transposable element presence using next-generation sequencing data. Nucleic Acids Research 39(6), 36 (2011). doi: 10.1093/nar/gkq1291

12. Linheiro, R.S., Bergman, C.M.: Whole genome resequencing reveals natural target site preferences of transposable elements in Drosophila melanogaster. PLoS One 7(2), 30008 (2012). doi: 10.1371/journal.pone.0030008. Accessed 2012-11-30.

13. Lee, E., Iskow, R., Yang, L., Gokcumen, O., Haseley, P., Luquette, L.J., Lohr, J.G., Harris, C.C., Ding, L., Wilson, R.K., Wheeler, D.a., Gibbs, R.a., Kucherlapati, R., Lee, C., Kharchenko, P.V., Park, P.J.: Landscape of somatic retrotransposition in human cancers. Science 337, 967–971 (2012). doi: 10.1126/science.1222077

14. Platzer, A., Nizhynska, V., Long, Q.: TE-Locate: a tool to locate and group transposable element occurrences using paired-end next-generation sequencing data. Biology 1(2), 395–410 (2012). doi: 10.3390/biology1020395

15. Kofler, R., Betancourt, A.J., Schlötterer, C.: Sequencing of pooled DNA samples (pool-seq) uncovers complex dynamics of transposable element insertions in Drosophila melanogaster. PLoS Genet 8(1), 1002487 (2012). doi: 10.1371/journal.pgen.1002487. Accessed 2012-11-29.

16. Nellaker, C., Keane, T.M., Yalcin, B., Wong, K., Agam, A., Belgard, T.G., Flint, J., Adams, D.J., Frankel, W.N., Ponting, C.P.: The genomic landscape shaped by selection on transposable elements across 18 mouse strains. Genome Biol 13(6), 45 (2012). doi: 10.1186/gb-2012-13-6-r45

17. Robb, S.M.C., Lu, L., Valencia, E., Burnette, J.M., Okumoto, Y., Wessler, S.R., Stajich, J.E.: The use of RelocaTE and unassembled short reads to produce high-resolution snapshots of transposable element generated diversity in rice. G3 3(6), 949–957 (2013). doi: 10.1534/g3.112.005348. Accessed 2013-07-10.

18. Chen, Y., Yao, H., Thompson, E.J., Tannir, N.M., Weinstein, J.N., Su, X.: VirusSeq: software to identify viruses and their integration sites using next-generation sequencing of human cancer tissue. Bioinformatics 29(2), 266–267 (2013). doi: 10.1093/bioinformatics/bts665. Accessed 2015-07-28.

19. Cridland, J.M., Macdonald, S.J., Long, A.D., Thornton, K.R.: Abundance and Distribution of Transposable Elements in Two Drosophila QTL Mapping Resources. Mol Biol Evol 30(10), 2311–2327 (2013). doi: 10.1093/molbev/mst129. Accessed 2015-06-25.

20. Zhuang, J., Wang, J., Theurkauf, W., Weng, Z.: TEMP: a computational method for analyzing transposable element polymorphism in populations. Nucleic Acids Res. 42(11), 6826–6838 (2014). doi: 10.1093/nar/gku323

21. Wu, J., Lee, W.-P., Ward, A., Walker, J.A., Konkel, M.K., Batzer, M.A., Marth, G.T.: Tangram: a comprehensive toolbox for mobile element insertion detection. BMC Genomics 15(1), 795 (2014). doi: 10.1186/1471-2164-15-795. Accessed 2015-06-25.

22. Nakagome, M., Solovieva, E., Takahashi, A., Yasue, H., Hirochika, H., Miyao, A.: Transposon Insertion Finder(TIF): a novel program for detection of de novo transpositions of transposable elements. BMC Bioinformatics 15, 71 (2014). doi: 10.1186/1471-2105-15-71

23. Gilly, A., Etcheverry, M., Madoui, M.-A., Guy, J., Quadrana, L., Alberti, A., Martin, A., Heitkam, T., Engelen, S., Labadie, K., Le Pen, J., Wincker, P., Colot, V., Aury, J.-M.: TE-Tracker: systematic identification of transposition events through whole-genome resequencing. BMC Bioinformatics 15, 377 (2014). doi: 10.1186/s12859-014-0377-z

24. Thung, D.T., Ligt, J.d., Vissers, L.E., Steehouwer, M., Kroon, M., Vries, P.d., Slagboom, E.P., Ye, K., Veltman, J.A., Hehir-Kwa, J.Y.: Mobster: accurate detection of mobile element insertions in next generation sequencing data. Genome Biology 15(10), 488 (2014). doi: 10.1186/s13059-014-0488-x. Accessed 2015-06-25.

25. Jiang, C., Chen, C., Huang, Z., Liu, R., Verdier, J.: ITIS, a bioinformatics tool for accurate identification of transposon insertion sites using next-generation sequencing data. BMC Bioinformatics 16(1), 72 (2015). doi: 10.1186/s12859-015-0507-2. Accessed 2015-04-29.

26. Fiston-Lavier, A.-S., Barrón, M.G., Petrov, D.A., González, J.: T-lex2: genotyping, frequency estimation and re-annotation of transposable elements using single or pooled next-generation sequencing data. Nucl. Acids Res. 43(4), 22–22 (2015). doi: 10.1093/nar/gku1250. Accessed 2016-12-14.

27. Hénaff, E., Zapata, L., Casacuberta, J.M., Ossowski, S.: Jitterbug: somatic and germline transposon insertion detection at single-nucleotide resolution. BMC Genomics 16, 768 (2015). doi: 10.1186/s12864-015-1975-5. Accessed 2016-12-13.

28. Rahman, R., Chirn, G.-w., Kanodia, A., Sytnikova, Y.A., Brembs, B., Bergman, C.M., Lau, N.C.: Unique transposon landscapes are pervasive across Drosophila melanogaster genomes. Nucl. Acids Res. 43(22), 10655–10672 (2015). doi: 10.1093/nar/gkv1193. Accessed 2016-12-13.

29. Hawkey, J., Hamidian, M., Wick, R.R., Edwards, D.J., Billman-Jacobe, H., Hall, R.M., Holt, K.E.: ISMapper: identifying transposase insertion sites in bacterial genomes from short read sequence data. BMC Genomics 16, 667 (2015). doi: 10.1186/s12864-015-1860-2. Accessed 2016-12-16.

30. Quadrana, L., Silveira, A.B., Mayhew, G.F., LeBlanc, C., Martienssen, R.A., Jeddeloh, J.A., Colot, V.: The Arabidopsis thaliana mobilome and its impact at the species level. eLife 5, 15716 (2016). doi: 10.7554/eLife.15716. Accessed 2016-12-13.

31. Kofler, R., Gómez-Sánchez, D., Schlötterer, C.: PoPoolationTE2: Comparative Population Genomics of Transposable Elements Using Pool-Seq. Mol Biol Evol 33(10), 2759–2764 (2016). doi: 10.1093/molbev/msw137. Accessed 2016-12-13.

32. Chen, J., Wrightsman, T., Wessler, S.R., Stajich, J.E.: RelocaTE2: a high resolution transposable element polymorphism mapping tool for population resequencing. Technical Report e2447v1, PeerJ Preprints (September 2016). https://peerj.com/preprints/2447 Accessed 2016-12-13.

33. Norel, R., Rice, J.J., Stolovitzky, G.: The self-assessment trap: can we all be better than average?Molecular Systems Biology 7(1) (2011). doi: 10.1038/msb.2011.70

34. Bergman, C.M.: A proposal for the reference-based annotation of de novo transposable element insertions. Mobile genetic elements 2(1), 51–54 (2012). doi: 10.4161/mge.19479

35. Rishishwar, L., Mariño-Ramírez, L., Jordan, I.K.: Benchmarking computational tools for polymorphic transposable element detection. Brief Bioinform, 072 (2016). doi: 10.1093/bib/bbw072. Accessed 2016-12-16.

36. Keane, T.M., Wong, K., Adams, D.J.: RetroSeq: transposable element discovery from next-generation sequencing data. Bioinformatics 29(3), 389–390 (2013). doi: 10.1093/bioinformatics/bts697. Accessed 2016-12-13.

37. Sudmant, P.H., Rausch, T., Gardner, E.J., Handsaker, R.E., Abyzov, A., Huddleston, J., Zhang, Y., Ye, K., Jun, G., Hsi-Yang Fritz, M., Konkel, M.K., Malhotra, A., Stutz, A.M., Shi, X., Paolo, Casale F., Chen, J., Hormozdiari, F., Dayama, G., Chen, K., Malig, M., Chaisson, M.J.P., Walter, K., Meiers, S., Kashin, S., Garrison, E., Auton, A., Lam, H.Y.K., Jasmine Mu, X., Alkan, C., Antaki, D., Bae, T., Cerveira, E., Chines, P., Chong, Z., Clarke, L., Dal, E., Ding, L., Emery, S., Fan, X., Gujral, M., Kahveci, F., Kidd, J.M., Kong, Y., Lameijer, E.-W., McCarthy, S., Flicek, P., Gibbs, R.A., Marth, G., Mason, C.E., Menelaou, A., Muzny, D.M., Nelson, B.J., Noor, A., Parrish, N.F., Pendleton, M., Quitadamo, A., Raeder, B., Schadt, E.E., Romanovitch, M., Schlattl, A., Sebra, R., Shabalin, A.A., Untergasser, A., Walker, J.A., Wang, M., Yu, F., Zhang, C., Zhang, J., Zheng-Bradley, X., Zhou, W., Zichner, T., Sebat, J., Batzer, M.A., McCarroll, S.A., The 1000 Genomes Project Consortium, Mills, R.E., Gerstein, M.B., Bashir, A., Stegle, O., Devine, S.E., Lee, C., Eichler, E.E., Korbel, J.O.: An integrated map of structural variation in 2,504 human genomes. Nature 526(7571), 75–81 (2015). doi: 10.1038/nature15394. Accessed 2016-12-16.

38. Stuart, T., Eichten, S., Cahn, J., Karpievitch, Y., Borevitz, J., Lister, R.: Population scale mapping of transposable element diversity reveals links to gene regulation and epigenomic variation. eLife 5, 20777 (2016). doi: 10.7554/eLife.20777. Accessed 2016-12-14.

39. Kaminker, J.S., Bergman, C.M., Kronmiller, B., Carlson, J., Svirskas, R., Patel, S., Frise, E., Wheeler, D.A., Lewis, S.E., Rubin, G.M., Ashburner, M., Celniker, S.E.: The transposable elements of the Drosophila melanogaster euchromatin: a genomics perspective. Genome Biol 3(12), 0084–1842 (2002). doi: 10.1186/gb-2002-3-12-research0084. Accessed 2016-12-13.

40. Goffeau, A., Barrell, B.G., Bussey, H., Davis, R.W., Dujon, B., Feldmann, H., Galibert, F., Hoheisel, J.D., Jacq, C., Johnston, M., Louis, E.J., Mewes, H.W., Murakami, Y., Philippsen, P., Tettelin, H., Oliver, S.G.: Life with 6000 genes. Science 274(5287), 546–5637 (1996)

41. Liti, G., Carter, D.M., Moses, A.M., Warringer, J., Parts, L., James, S.A., Davey, R.P., Roberts, I.N., Burt, A., Koufopanou, V., Tsai, I.J., Bergman, C.M., Bensasson, D., O’Kelly, M.J.T., Oudenaarden, A.v., Barton, D.B.H., Bailes, E., Nguyen, A.N., Jones, M., Quail, M.A., Goodhead, I., Sims, S., Smith, F., Blomberg, A., Durbin, R., Louis, E.J.: Population genomics of domestic and wild yeasts. Nature 458(7236), 337–41 (2009). doi: 10.1038/nature07743

42. Strope, P.K., Skelly, D.A., Kozmin, S.G., Mahadevan, G., Stone, E.A., Magwene, P.M., Dietrich, F.S., McCusker, J.H.: The 100-genomes strains, an S. cerevisiae resource that illuminates its natural phenotypic and genotypic variation and emergence as an opportunistic pathogen. Genome Res. 25(5), 762–774 (2015). doi: 10.1101/gr.185538.114. Accessed 2015-10-22.

43. Almeida, P., Barbosa, R., Zalar, P., Imanishi, Y., Shimizu, K., Turchetti, B., Legras, J.-L., Serra, M., Dequin, S., Couloux, A., Guy, J., Bensasson, D., Goncalves, P., Sampaio, J.P.: A Population Genomics Insight into the Mediterranean Origins of Wine Yeast Domestication. Mol. Ecol. (2015). doi: 10.1111/mec.13341

44. Kim, J.M., Vanguri, S., Boeke, J.D., Gabriel, A., Voytas, D.F.: Transposable elements and genome organization: a comprehensive survey of retrotransposons revealed by the complete Saccharomyces cerevisiae genome sequence. Genome Research 8(5), 464–78 (1998)

45. Carr, M., Bensasson, D., Bergman, C.M.: Evolutionary genomics of transposable elements in Saccharomyces cerevisiae. PLoS ONE 7(11), 50978 (2012). doi: 10.1371/journal.pone.0050978. Accessed 2012-12-03.

46. Gafner, J., Philippsen, P.: The yeast transposon Ty1 generates duplications of target DNA on insertion. Nature 286(5771), 414–418 (1980). doi: 10.1038/286414a0. Accessed 2015-08-27.

47. Ji, H., Moore, D.P., Blomberg, M.A., Braiterman, L.T., Voytas, D.F., Natsoulis, G., Boeke, J.D.: Hotspots for unselected Ty1 transposition events on yeast chromosome III are near tRNA genes and LTR sequences. Cell 73 (5), 1007–1018 (1993). doi: 10.1016/0092-8674(93)90278-X. Accessed 2015-07-20.

48. Chalker, D.L., Sandmeyer, S.B.: Transfer RNA genes are genomic targets for de Novo transposition of the yeast retrotransposon Ty3. Genetics 126(4), 837–850 (1990). Accessed 2016-12-13.

49. Chalker, D.L., Sandmeyer, S.B.: Ty3 integrates within the region of RNA polymerase III transcription initiation. Genes Dev. 6(1), 117–128 (1992). doi: 10.1101/gad.6.1.117. Accessed 2016-12-13.

50. Devine, S.E., Boeke, J.D.: Integration of the yeast retrotransposon Ty1 is targeted to regions upstream of genes transcribed by RNA polymerase III. Genes Dev. 10(5), 620–633 (1996). doi: 10.1101/gad.10.5.620. Accessed 2016-12-13.

51. Rinckel, L.A., Garfinkel, D.J.: Influences of histone stoichiometry on the target site preference of retrotransposons Ty1 and Ty2 in Saccharomyces cerevisiae. Genetics 142(3), 761–776 (1996)

52. Zou, S., Ke, N., Kim, J.M., Voytas, D.F.: The Saccharomyces retrotransposon Ty5 integrates preferentially into regions of silent chromatin at the telomeres and mating loci. Genes Dev. 10(5), 634–645 (1996)

53. Baller, J.a., Gao, J., Stamenova, R., Curcio, M.J., Voytas, D.F.: A nucleosomal surface defines an integration hotspot for the Saccharomyces cerevisiae Ty1 retrotransposon. Genome Research 22(4), 704–13 (2012). doi: 10.1101/gr.129585.111

54. Mularoni, L., Zhou, Y., Bowen, T., Gangadharan, S., Wheelan, S.J., Boeke, J.D.: Retrotransposon Ty1 integration targets specifically positioned asymmetric nucleosomal DNA segments in tRNA hotspots. Genome research 22(4), 693–703 (2012). doi: 10.1101/gr.129460.111

55. Qi, X., Daily, K., Nguyen, K., Wang, H., Mayhew, D., Rigor, P., Forouzan, S., Johnston, M., Mitra, R.D., Baldi, P., Sandmeyer, S.: Retrotransposon profiling of RNA polymerase III initiation sites. Genome Research 22(4), 681–92 (2012). doi: 10.1101/gr.131219.111

56. Baller, J.A., Gao, J., Voytas, D.F.: Access to DNA establishes a secondary target site bias for the yeast retrotransposon Ty5. Proceedings of the National Academy of Sciences of the United States of America 108(51), 20351–20356 (2011). doi: 10.1073/pnas.1103665108

57. Zou, S., Wright, D.A., Voytas, D.F.: The Saccharomyces Ty5 retrotransposon family is associated with origins of DNA replication at the telomeres and the silent mating locus HMR. PNAS 92(3), 920–924 (1995). Accessed 2016-12-13.

58. Li, H., Handsaker, B., Wysoker, A., Fennell, T., Ruan, J., Homer, N., Marth, G., Abecasis, G., Durbin, R., Subgroup, .G.P.D.P.: The Sequence Alignment/Map format and SAMtools. Bioinformatics 25(16), 2078–2079 (2009). doi: 10.1093/bioinformatics/btp352. Accessed 2016-12-13.

59. Lee, H., Schatz, M.C.: Genomic dark matter: The reliability of short read mapping illustrated by the genome mappability score. Bioinformatics 28(16), 2097–2105 (2012). doi: 10.1093/bioinformatics/bts330

60. Quinlan, A.R., Hall, I.M.: BEDTools: a flexible suite of utilities for comparing genomic features. Bioinformatics 26(6), 841–842 (2010). doi: 10.1093/bioinformatics/btq033. Accessed 2015-02-18.

61. Kent, W.J., Zweig, A.S., Barber, G., Hinrichs, A.S., Karolchik, D.: BigWig and BigBed: enabling browsing of large distributed datasets. Bioinformatics 26(17), 2204–2207 (2010). doi: 10.1093/bioinformatics/btq351. Accessed 2015-09-15.

62. Stempor, P.: Seqplots: An Interactive Tool for Visualizing NGS Signals and Sequence Motif Densities Along Genomic Features Using Average Plots and Heatmaps., (2014). R package version 1.6.0. http://github.com/przemol/seqplots

63. Bardou, P., Mariette, J., Escudié, F., Djemiel, C., Klopp, C.: jvenn: an interactive Venn diagram viewer. BMC Bioinformatics 15(1), 293(2014). doi: 10.1186/1471-2105-15-293. Accessed 2015-10-22.

64. Team, R.D.C.: R: a Language and Environment for Statistical Computing, Vienna, Austria (2012). ISBN 3-900051-07-0. http://www.R-project.org/

65. Stajich, J.E., Block, D., Boulez, K., Brenner, S.E., Chervitz, S.A., Dagdigian, C., Fuellen, G., Gilbert, J.G.R., Korf, I., Lapp, H., Lehväslaiho, H., Matsalla, C., Mungall, C.J., Osborne, B.I., Pocock, M.R., Schattner, P., Senger, M., Stein, L.D., Stupka, E., Wilkinson, M.D., Birney, E.: The Bioperl Toolkit: Perl Modules for the Life Sciences. Genome Res. 12(10), 1611–1618 (2002). doi: 10.1101/gr.361602. Accessed 2016-12-13.

66. Smit, A.F.A., Hubley, R., Green, P.: RepeatMasker, (2013). http://www.repeatmasker.org

67. Kuhn, R.M., Haussler, D., Kent, W.J.: The UCSC genome browser and associated tools. Briefings in Bioinformatics, 1–18 (2012). doi: 10.1093/bib/bbs038

68. Kent, W.J.: BLAT—The BLAST-Like Alignment Tool. Genome Res. 12(4), 656–664 (2002). doi: 10.1101/gr.229202. Accessed 2013-02-25.

69. Slater, G.S.C., Birney, E.: Automated generation of heuristics for biological sequence comparison. BMC Bioinformatics 6, 31 (2005). doi: 10.1186/1471-2105-6-31

70. Langmead, B., Trapnell, C., Pop, M., Salzberg, S.L.: Ultrafast and memory-efficient alignment of short DNA sequences to the human genome. Genome Biology 10(3), 25(2009). doi: 10.1186/gb-2009-10-3-r25. Accessed 2015-09-15.

71. Li, H.: Aligning sequence reads, clone sequences and assembly contigs with BWA-MEM. arXiv, 1303–3997 (2013). Accessed 2015-02-18.

